# RD-OMICS: An Integrative Multi-Omics Data Inventory in Rare Diseases

**DOI:** 10.64898/2026.06.29.735296

**Authors:** Huanfei Wang, Shixue Sun, Ewy A. Mathé, Qian Zhu

## Abstract

Rare diseases (RD) impact over 30 million individuals in the United States, yet fewer than 5% of the identified conditions have FDA-approved treatments. Progress in RD research is hindered by small patient cohorts, biological heterogeneity, and the fragmented, inconsistently annotated publicly available omics data, which limits integrative analysis and translational discovery.

Here, we present RD-OMICS, a data inventory with integrated and structured RD omics data from Gene Expression Omnibus (GEO), in the form of a knowledge graph. We developed a metadata harmonization pipeline that combines rule-based mapping and large language model (LLM)-assisted semantic categorization. The graph-based data model was defined to integrate different types of data including disease conditions, experiments, samples, platforms, projects, and publications into a centralized inventory graph.

In this preliminary study, 11,049 GEO series for 126 rare diseases were processed and integrated into RD-OMICS, which includes 375,930 individual biospecimen samples, 1,578 sequencing and array platforms, 10,938 biological projects. Case studies demonstrate the use of RD-OMICS in supporting rare disease research, omics cohort construction, and transcriptome-based drug repurposing for amyotrophic lateral sclerosis (ALS).

RD-OMICS provides a scalable foundation for transforming fragmented omics data into a structured, harmonized and interoperable resource, facilitating therapeutic development and other translational discoveries in rare diseases.

## Introduction

Rare diseases (RD) affect over 30 million individuals in the United States and approximately 300 million worldwide, yet only about 500 of the more than 10,000 known rare conditions have FDA-approved treatments (1–3). Research in this area remains challenging due to small patient populations, genetic heterogeneity, and limited availability of well-characterized datasets (4). These constraints impede both mechanistic understanding of RD and the development of effective diagnostics and therapeutics. At the same time, data-driven approaches that supports integration and interpretation of heterogeneous biological data are increasingly being explored in the RD filed.

Omics data offers comprehensive characterization and quantification of biological molecules, such as DNA, RNA, proteins, epigenetic modifications, and metabolites that collectively define the structure, function, and dynamic states of biological systems (5). Advances in omics technologies and integrative bioinformatics approaches have transformed translational research by supporting systematic interrogation of disease mechanisms, biomarker discovery, and therapeutic target identification (6, 7). Across diverse disease contexts, including cancer and neurological disorders, integrative analysis of omics datasets has provided critical insights into molecular dysregulation that are not accessible through single-layer analyses alone (8–10).

Omics-based biomedical research plays a particularly critical role in RD research. Genomic and transcriptomic analyses have substantially improved multiple aspects of RD healthcare by enabling molecular diagnosis, disease subtyping, and identification of pathogenic mechanisms that might otherwise remain unresolved (11, 12). Integrative analysis of multi-omics data provides powerful frameworks for uncovering dysregulated pathways and molecular signatures that can support therapeutic discovery and drug repurposing (13–15). By leveraging disease-associated gene expression profiles, omics-driven approaches are increasingly being applied to prioritize candidate drugs for RDs, offering a cost-effective and time-efficient alternative to de novo drug development (16, 17).

Major advances in omics technologies, together with the availability of public data repositories such as the Gene Expression Omnibus (GEO) (18, 19), have created valuable opportunities for secondary analysis and integrative analysis of omics studies in RDs (20–22). As one of the largest public repositories for high-throughput functional genomics data, GEO contains microarray and next-generation sequencing datasets evaluating in vitro, in vivo, or human biological samples spanning thousands of diseases, including RD, and therefore represents an important resource for RD research (23). However, despite its broad coverage and scientific value, GEO data are not immediately ready for systematic reuse. Because datasets are submitted by individual users, metadata quality, terminology, sample annotations, study descriptions, and experimental details can vary substantially across submissions (24, 25). As a result, researchers must devote considerable time and effort to identify relevant datasets, interpret submitter provided metadata, harmonize sample-level information across studies, and format data into analysis ready formats. This challenge is particularly important in RD research, where limited and fragmented omics datasets must be efficiently identified, linked, and integrated across studies to enable meaningful secondary analyses.

To address these challenges, we present RD-OMICS, a data inventory comprising harmonized experiment-and sample-level metadata from GEO for a collection of RDs. RD-OMICS organizes datasets into a graph-based data model that links conditions, projects, experiments, samples, platforms, and publications, thereby enabling scalable, disease-centric integration and systematic exploration of omics data. Through demonstrated case studies, we show how RD-OMICS can support efficient dataset discovery, cohort construction, and downstream analytical applications in RD research. In this pilot study, we focus on GEO data, with planned extensions to additional repositories, including ArrayExpress (26), PRIDE (27), and MetaboLights (28), to enable broader omics integration in future work.

## Materials

GEO served as the primary data source for this study. Multiple GEO sections capture complementary information that is useful for omics data discovery and secondary analysis. GEO Series (GSE) describe study-level information, including overall experimental design, research objective, and associated datasets. GEO Sample (GSM) provides sample-level metadata, such as biological source, disease or condition, tissue or cell type, treatment status, and other sample characteristics. GEO Platform (GPL) describes the technologies and platforms used to generate the data, including microarray or sequencing platforms. BioProject (PRJNA) provides project-level information that links related sequencing datasets and supports broader contextualization of the study. More details about these sections can be found in Table 1.

**Table 1.** GEO data description.

| Sections | Description |
| --- | --- |
| GEO Series (GSE) | It defines the overall study context and links together related samples. These entries include study summaries, data tables, and experimental annotations. |
| BioProjects (PRJNA) | It consolidates datasets generated by a single institution or consortium under a shared research initiative [4], and provide project-level metadata such as title, description, and organizational affiliation. |
| GEO Sample (GSM) | A Sample entry describes the biological material used in an experiment, their treatment conditions, and the molecular abundance measurements derived from each. Each sample is linked to one Platform and may participate in multiple Series. |
| GEO Platform (GPL) | A Platform entry specifies the array or sequencing technology used to generate the data. For array-based platforms, it also includes probe annotations or template definitions. A single Platform may be referenced by many different Samples. |

## Methods

To develop RD-OMICS, we identified, extracted, and harmonized rare disease–related omics data from GEO and represented it in the form of a knowledge graph. The overall workflow included four main components: rare disease selection, GEO data retrieval, data normalization and harmonization, and RD-OMICS development, shown in Figure 1. Implementation details about each component are summarized in the following sections.

**Figure 1.**
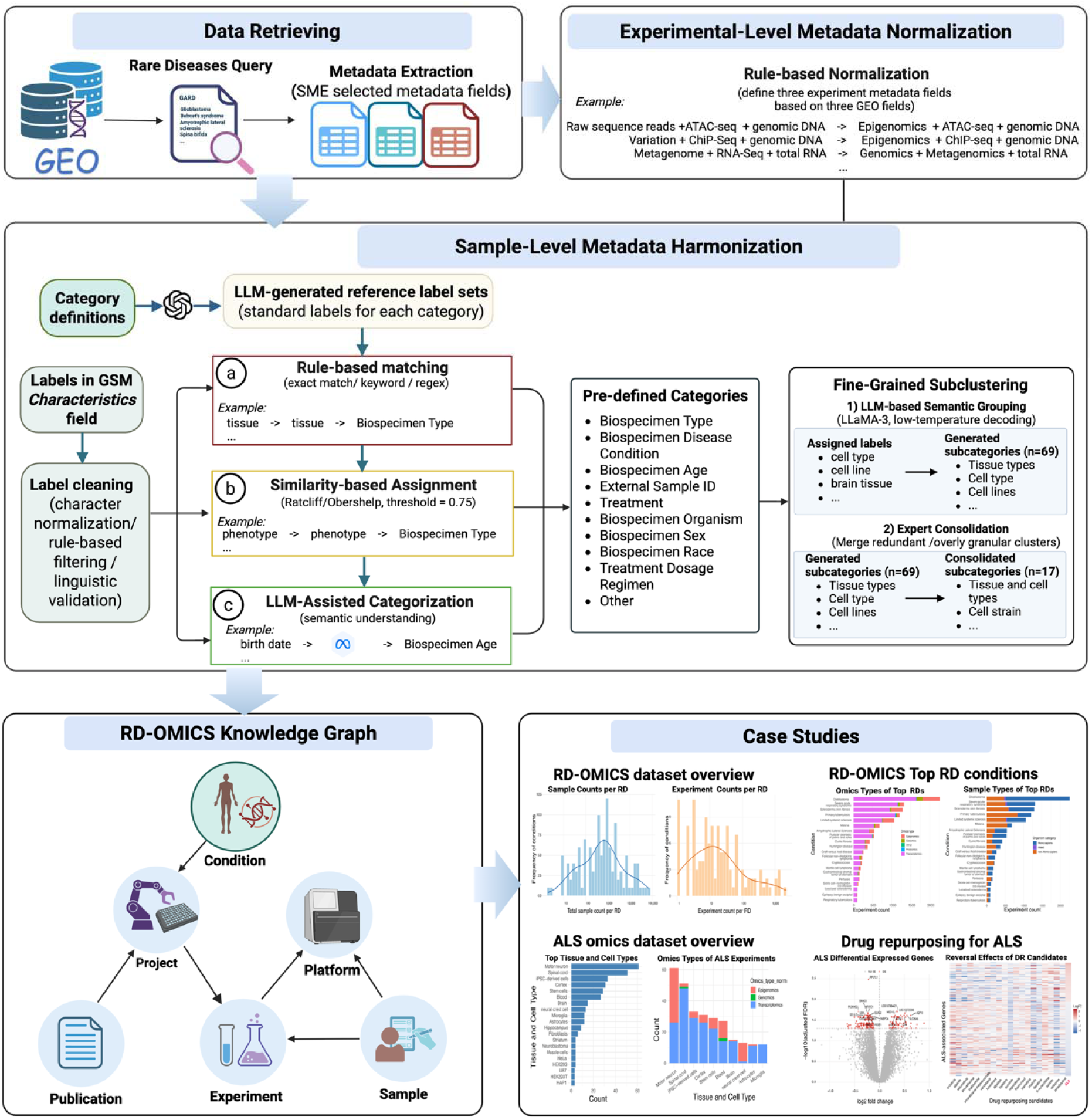
Overview of the RD-OMICS framework. RD–associated omics data are retrieved from GEO and processed through experiment-level and sample-level metadata harmonization pipelines. The resulting harmonized metadata are integrated into the RD-OMICS knowledge graph. Created in BioRender. Sun, S. (2026) https://BioRender.com/7u37az1.

## 1. RD List Generation

As a pilot study, we curated a list of RDs using three selection criteria. First, we included Glioblastoma multiforme (GBM)-related RDs, motivated by the successful use of GEO derived omics data for drug repurposing in GBM (16), with the goal of extending this prior approach to a broader set of RDs. We selected 92 RDs with strong genetic associations to GBM, as identified through our previous RD clustering analysis (29).

Second, we included RDs with relatively high prevalence. The *All of Us* Research Program (30) aggregates diverse data types from participants across the United States, including electronic health record (EHR) and genomics data, enabling large-scale, integrative biomedical analyses. We mapped over 10,000 RDs obtained from the National Center for advancing Translational Sciences (NCATS) Genetic and Rare Diseases (GARD) Information Center (31) to the *All of Us* and selected the top 100 RDs with the highest observed prevalence.

Third, we collected RDs of specific research interest. Five additional RDs were manually selected based on ongoing research priorities: Juvenile Neuronal Ceroid Lipofuscinosis (JNCL, also known as CLN3 disease), Niemann-Pick disease type C (NPC), Scleroderma skin fibrosis, Prader Willi-syndrome, and Amyotrophic lateral sclerosis (ALS).

After removing duplicates, a final set of 194 distinct RDs was obtained. Disease information, including GARD IDs, preferred disease names and disease synonyms, was extracted from GARD and used as the foundation for RD-OMICS development. The entire list of selected RDs is provided in the Supplementary File 1 named “RD_List.xlsx”.

## 2. Data Extraction from GEO

Omics data was extracted from GEO using a two-step approach: (1) identification of relevant GSEs for each selected RD and (2) for each GSE, data retrieval from different sections, including GSM, GPL, and PRJNA.

To identify disease-associated GEO Series, we queried GARD disease names and synonyms via the NCBI Entrez search API (32). The resulting GSE accession IDs were used as entry points for subsequent data extraction. Because NCBI API does not provide all experiment-and sample-level data elements for downstream analysis, we developed a scripted extraction workflow in Python with BeautifulSoup (33) to retrieve data from GSE, GSM, GPL, and PRJNA sections. To improve scalability, data retrieval was parallelized across accessions, while robustness was ensured through timeout control, automatic retries, and logging of failed requests for targeted reprocessing.

Metadata fields within these GEO sections were reviewed to determine their relevance to characterization of projects, experiments, samples, platforms, and publications. Table 2 summarizes the data fields extracted from corresponding GEO sections.

**Table 2.** Description about the metadata fields extracted from GEO.

| GEO Sections | Extracted Metadata Fields |
| --- | --- |
| GEO Series (GSE) | Title, summary, linked samples and platforms, associated BioProject ID, supplementary file links, and PubMed IDs and references when available. |
| BioProjects (PRJNA) | Project title, data type, summary of project, and submission metadata (date, submitter, affiliations, etc.). |
| GEO Sample (GSM) | Organism, sample characteristics (biospecimen type, cell line, treatment, genotype, or donor properties (e.g., age, sex), sample process, molecular extracted molecular, etc. |
| GEO Platform (GPL) | Platform name, manufacturer, and sequencing or array design specifications. |

## 3. Data Normalization and Harmonization

Although standardized guidelines for GEO data submission are available (34), GEO datasets remain difficult to reuse systematically because experiment-and sample-level data are often inconsistently labeled, variably structured, and incompletely standardized. As a result, researchers must often perform extensive manual review and curation before GEO datasets can be discovered, filtered, integrated, or analyzed for secondary analysis (24). Thus, we developed a structured framework for harmonizing and integrating GEO-derived rare disease omics data towards RD-OMICS development.

### 3.1 Experiment-level Data Normalization

Experiment-level metadata is distributed across multiple GEO sections and fields, while no single field adequately captures all essential aspects of an omics experiment. For example, the *Data type* field may describe a study broadly as “Transcriptome or Gene expression”, but this description does not clearly specify the measured molecular layer, sequencing or assay strategy, or input molecular material. To obtain a complete set of experiment-level metadata, we adopted a rule-based normalization approach that leverages information from multiple experiment-related metadata fields.

We manually selected three experiment-related fields including *Data type*, *Library strategy*, and *Extracted molecule* across GSE, PRJNA, and GSM sections, respectively. We normalized the data from these selected fields into three experiment-level attributes: omics domain, which represents the biological data modality; assay type, which represents the sequencing or assay strategy; and extracted molecule, which represents the input molecular material used for library preparation or measurement. The normalization process was performed by leveraging the controlled vocabularies including the ontology of data analysis and management (EDAM) (35), National Cancer Institute Thesaurus (NCTI) (36), Ontology of RNA Sequencing (ORNASEQ) (37), Sequence Types and Features Ontology (SO) (38). The complete lists of normalization rules and terminology sources are provided in Supplementary File 2 (“Experiment_metadata_normalization_rules.xlsx”) and File 3 (“Term_definition_source.docx”). Representative examples of normalized experiment-level data are shown in Table 3.

**Table 3.**
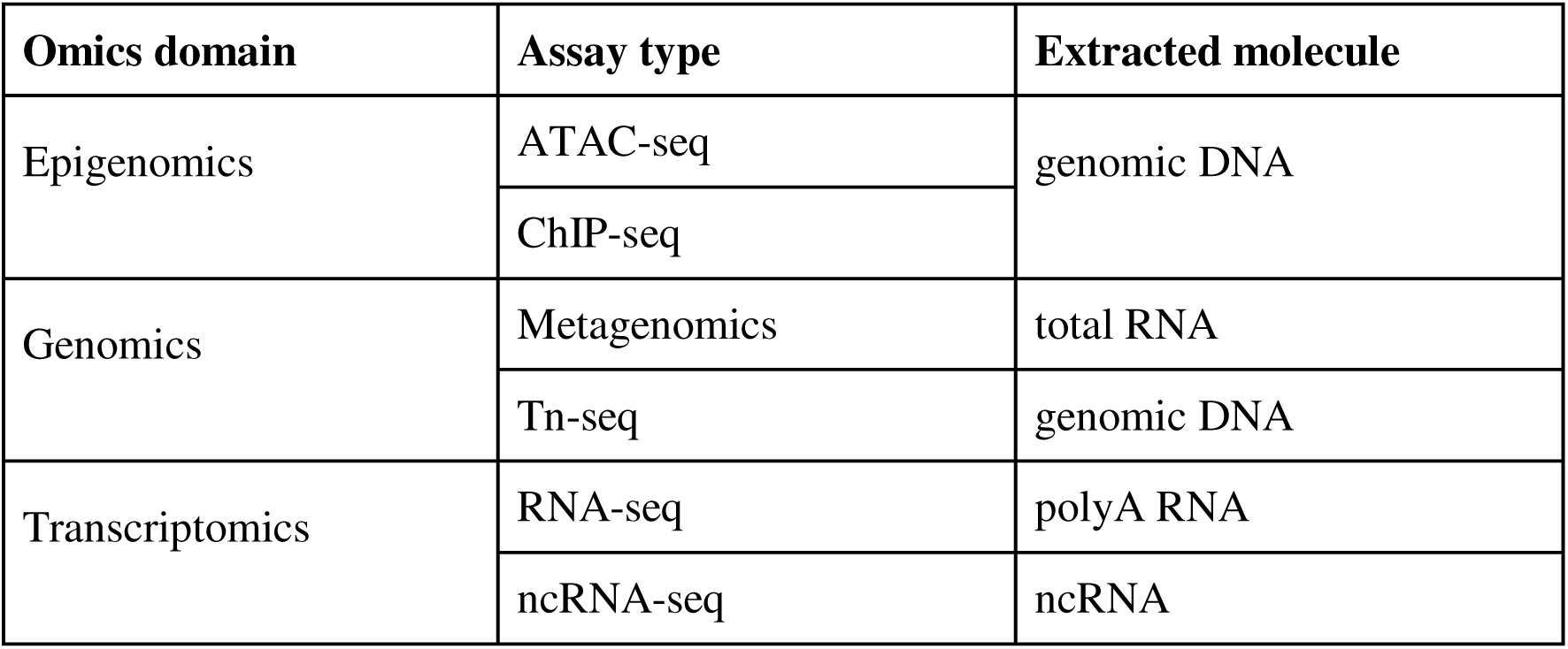
Examples of normalized experiment-level metadata drawn from GSE, PRJNA, and GSM sections.

| Omics domain | Assay type | Extracted molecule |
| --- | --- | --- |
| Epigenomics | ATAC-seq | genomic DNA |
|  | ChIP-seq |  |
| Genomics | Metagenomics | total RNA |
|  | Tn-seq | genomic DNA |
| Transcriptomics | RNA-seq | polyA RNA |
|  | ncRNA-seq | ncRNA |

### 3.2 Sample-level Data Normalization and Harmonization

To enable standardized sample-level data representation and consistent comparison of biospecimens across studies, we normalized data from the GSM *Characteristics* field. The *Characteristics* field contains sample data as label–value pairs in free-text. These labels and their associated values describe biospecimen-related information, such as tissue type, cell line, treatment, genotype. However, these labels vary substantially across studies in wording, specificity, and representation format, which make it difficult to compare and integrate sample-level metadata across GEO datasets. For example, the *Characteristics* entry for GSM8655443 is “tissue: whole body; genotype: WT; time: 45 hpf,” which includes the labels *tissue*, *genotype*, and *time*. By comparison, GSM1538996 contains “genotype: WBS; donor: GDB-306; gender: male; culture: Single-cell culture; spike-in: ERCC ExFold mix 2; library-type: unstranded 100bp PE reads,” Thus, we developed a two-tiered harmonization strategy for sample-level metadata. First, labels were assigned to ten self-defined biomedical categories based on rule-based matching, string-similarity scoring, and large language model (LLM)-assisted classification. Second, labels within each mapped category were further organized into fine-grained subcategories using LLM-based semantic grouping. The resulting standardized categories and subcategories were encoded as structured sample-level data properties in the RD-OMICS.

#### 3.2.1 *Characteristics* Label Cleaning

We obtained and cleaned a unique list of labels from the *Characteristics* field. We removed extraneous punctuation, unmatched brackets, trailing quotation marks, excessively long labels, and labels composed only of numbers. We also excluded invalid or non-informative labels, such as file paths, identifiers, or ambiguous short codes (e.g., #25, Tg(-7.2sox10), /usr/bin/). Furthermore, we performed linguistic validation by tokenizing compound labels and cross-referencing the resulting tokens with the Natural Language Toolkit (NLTK) (39). Labels composed primarily of non-English strings, random codes, or technical artifacts were removed.

#### 3.2.2 Semantic Categorization

To enable harmonization of the retrieved sample-level metadata, we predefined ten biomedical categories that could capture common attributes, which was informed by prior studies (25, 40) emphasizing the importance of data completeness and standardization for data reuse and reproducibility. The category definitions were aligned with the biomedical vocabularies and ontologies, including the Biological Collections Ontology (BCO) (41), General Formal Ontology for Biology (GFI-BIO) (42), and National Cancer Institute Thesaurus (NCIT) (36) (Supplementary File 3, “Term_definition_source.docx”). Table 4 summarizes the ten predefined biomedical categories and their definitions, including *Biospecimen Type*, *Biospecimen Disease Condition*, *Biospecimen Age*, *External Sample ID*, *Treatment*, *Biospecimen Organism*, *Biospecimen Sex*, *Biospecimen Race*, *Treatment Dosage Regimen*, and *Other*. We then mapped the sample-level labels to the ten categories via (a) rule-based match, (b) string similarity comparison, and (c) LLM-assisted categorization.

**Table 4.**
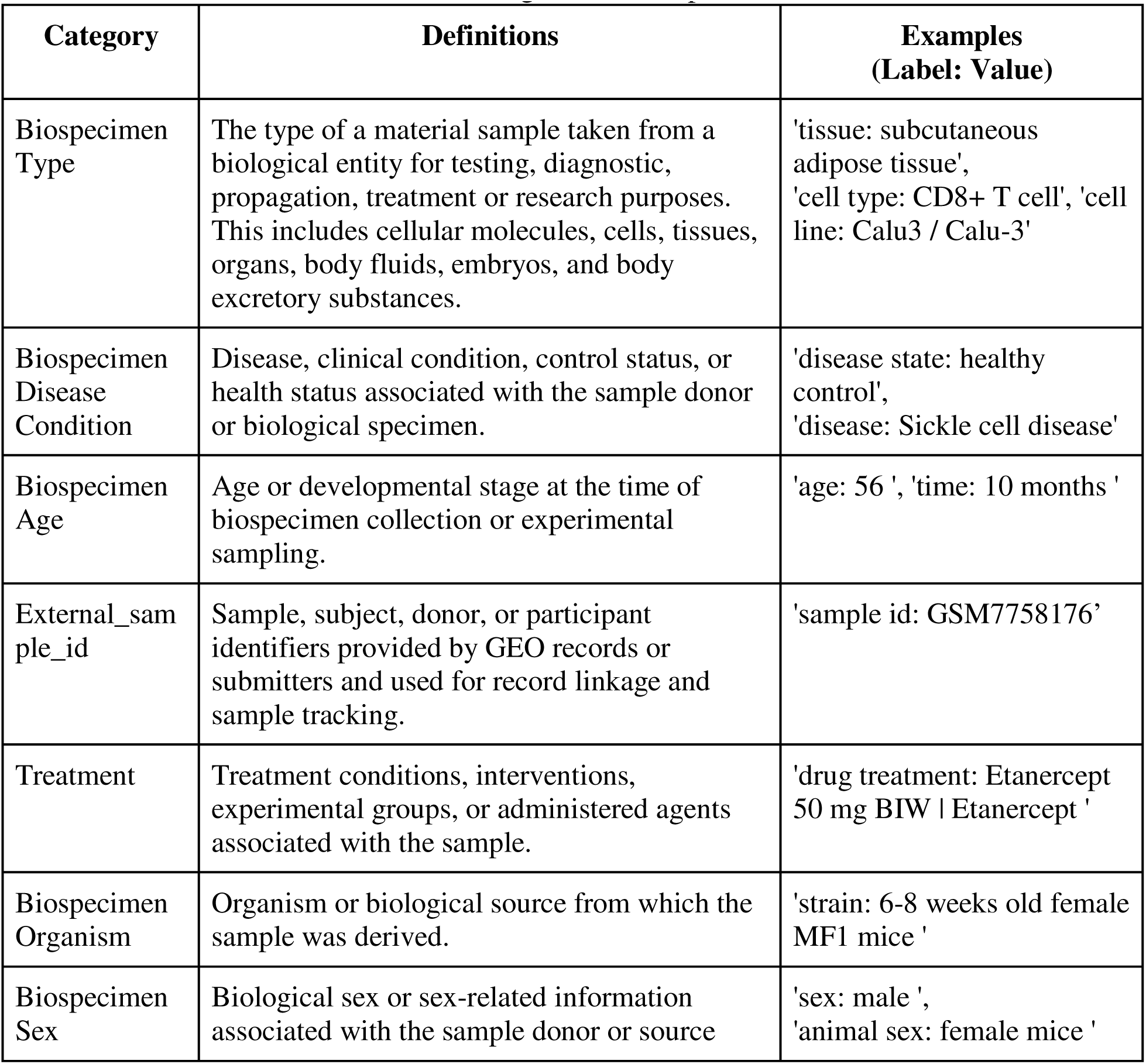
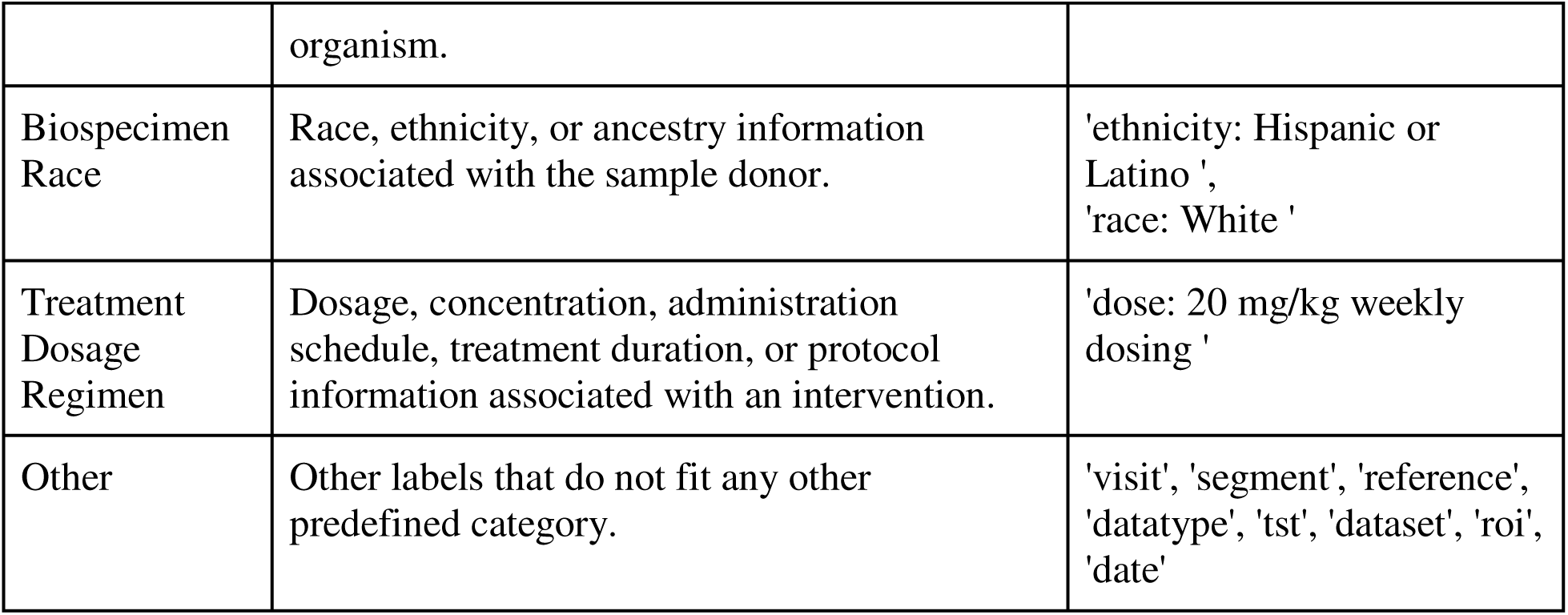
Pre-defined biomedical categories for sample-level metadata harmonization.

### a) Rule-based Match

Based on the predefined categories (Table 4), we first compiled category reference information, including category names, definitions, synonyms, and representative examples (Supplementary File 4 “Category_reference_information.csv”). This predefined category was then provided to the model GPT-5.3 (43) to generate an enhanced category-specific mapping dictionary for standardizing heterogeneous GEO labels. For each category, the GPT was instructed to generate a dictionary that includes three types of matching rules: exact-match terms, keyword-match terms, and regular-expression patterns designed to capture common label structures and naming variants. This dictionary served as the target vocabulary for standardizing heterogeneous GEO labels into predefined categories. A prompt example is provided in Supplementary File 5 (“Semantic_categorization_sample_characteristics.xlsx”).

For each raw GEO label, we first performed exact string matching against the exact-match terms in the dictionary. Raw labels were assigned to this category if they matched a dictionary term. Labels that could not be assigned by exact match were then evaluated using keyword-based matching. In this step, the raw GEO labels were compared with the GPT-generated category-specific keywords, which captured semantically related terms and common naming variations for each predefined category. For the remaining unmapped labels, regular-expression patterns were applied to capture common GEO label structures and naming variants, such as labels ending in id or labels containing terms related to tissue, age, disease, treatment, dose, or sample source. All rule-based assignments were restricted to the predefined category set, and no new categories were introduced during this step. Table 5 shows examples of rule-based category match.

**Table 5.**
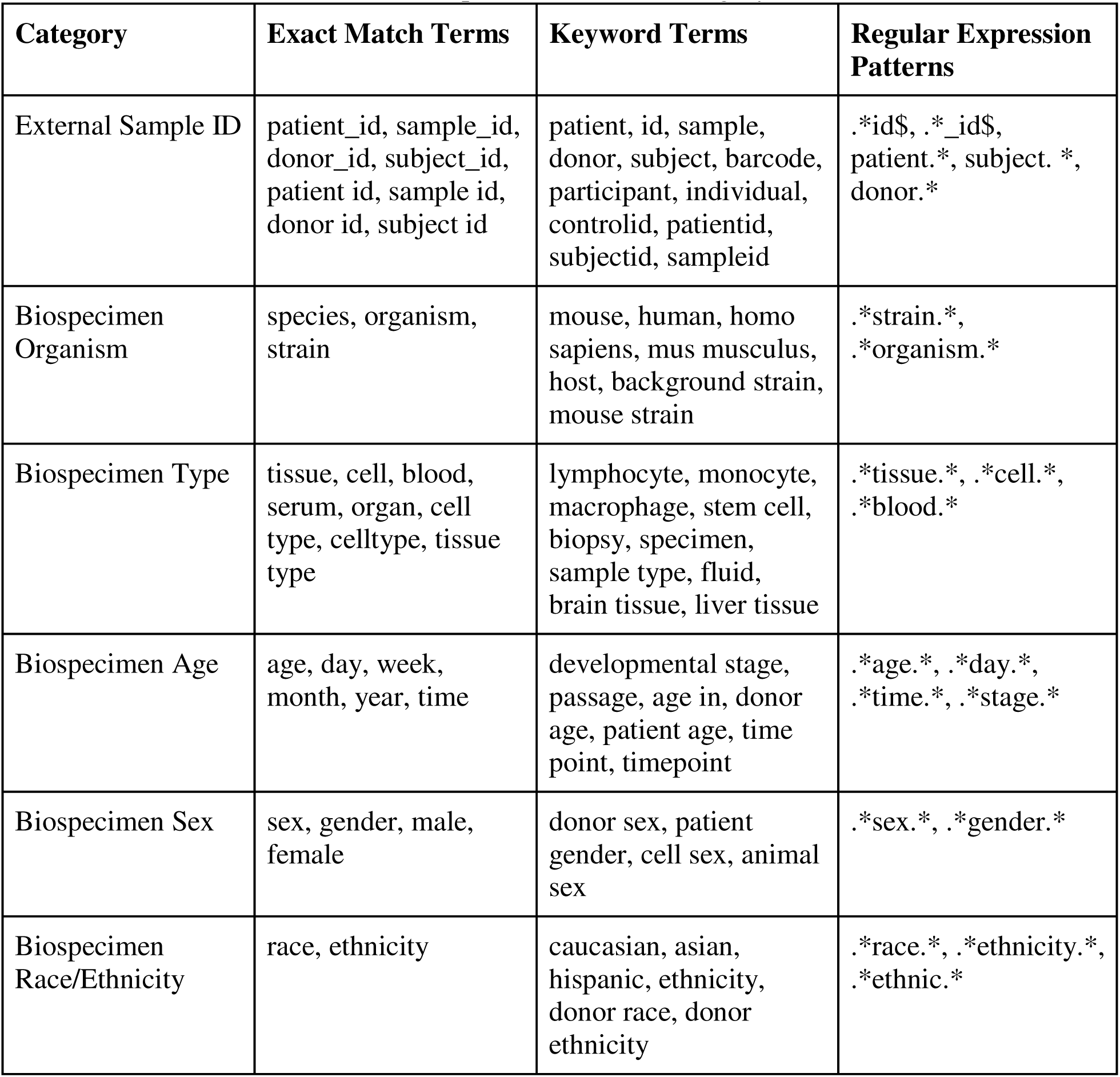
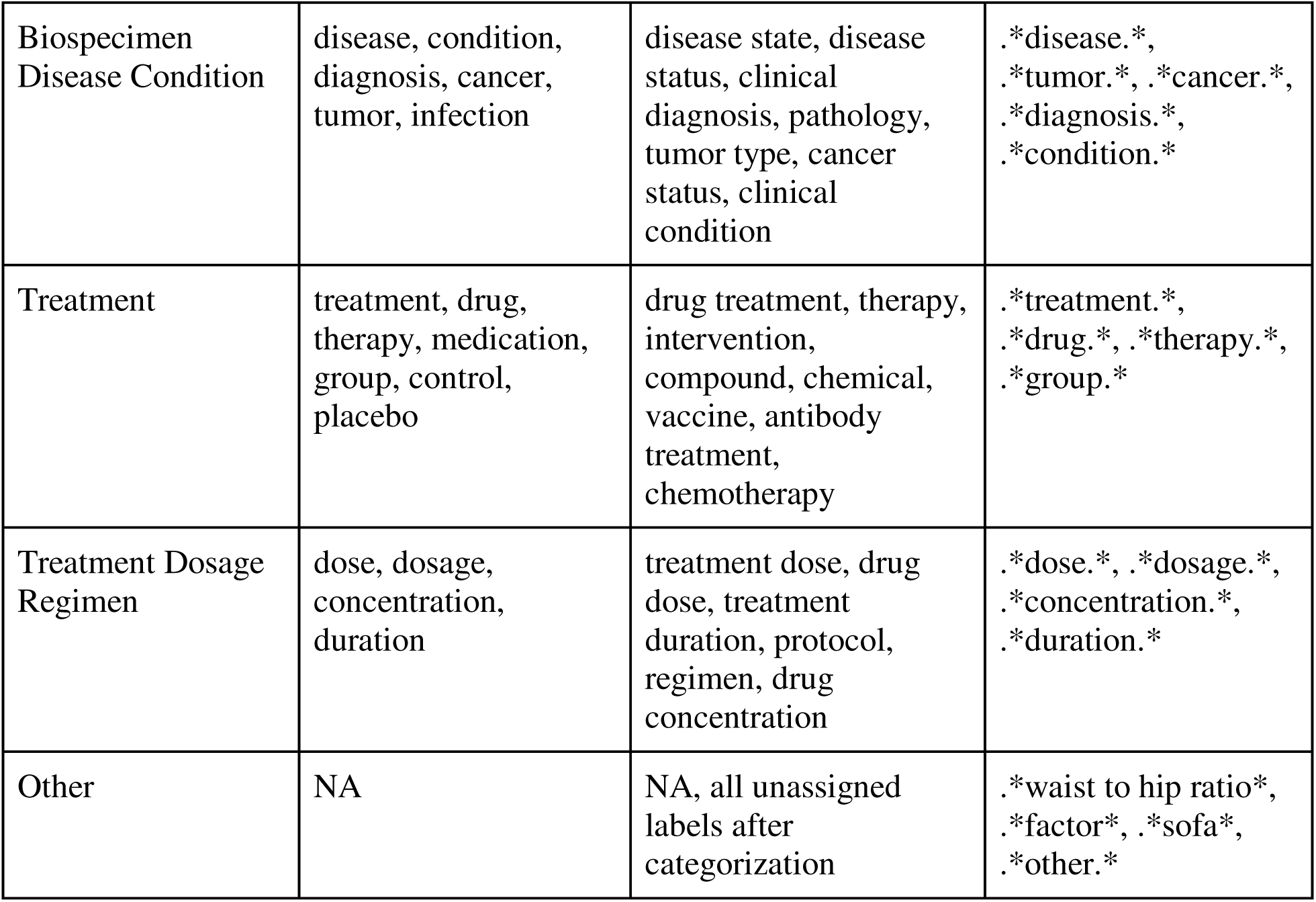
Examples of rule-based category match.

### b) String Similarity-based Match

Labels that remained unassigned were then evaluated using string-similarity-based match. This step was aimed to capture labels that were lexically similar to labels already assigned in the rule-based step (as reference labels). Each unmapped label was compared with reference labels using Python’s ‘difflib.SequenceMatcher’, which outputs a character-level similarity score from 0 to 1. Higher scores indicate greater similarity between label names for category assignment. The reference label with the highest similarity score was selected as the best match. The cutoff of 0.75 was selected after preliminary testing of alternative thresholds to balance label coverage and assignment accuracy. If the best-match score was ≥ 0.75, the unassigned label was assigned to the same category as the matched reference label. If the best-match score was < 0.75, the label remained unassigned and was passed to the subsequent LLM-based assignment step.

### c) LLM-Assisted Categorization

Labels that remained unassigned after steps (a) and (b) were categorized using the Llama-3.1-70B-Instruct model (44). For each LLM query, the prompt was provided with semantic context, such as category names, definitions, predefined label vocabulary, and synonyms for mapping those unassigned labels to the predefined biomedical categories. The model was instructed to assign each label to the most appropriate predefined category, use ‘Other’ when no category fit well, and return only structured JSON containing the raw label, assigned category, confidence level, and a brief rationale. Labels were sorted by occurrence count and processed in batches of eight to keep prompts concise while preserving sufficient context for consistent categorization. A low temperature setting of 0.02 was used to reduce output variability.

After generation, the JSON response was programmatically parsed and validated against the controlled category list. New categories were not allowed. If the model returned a category outside the predefined biomedical vocabulary, that label was reassigned to ‘Other’ and marked as low confidence. If a batch did not return valid JSON, the batch was retried using smaller subsets; labels from failed batches were assigned to ‘Other’. Examples of LLM-based label categorization are listed in Table 6, and the full list of assigned labels after the three steps are provided in Supplementary File 5 (“Semantic_categorization_sample_characteristics. xlsx”).

**Table 6.** Examples of LLM-assisted label categorization.

| Raw Label | Assigned Category | Confidence | LLM Reasoning |
| --- | --- | --- | --- |
| birth date | Biospecimen Age | HIGH | Birth date is used to calculate age, which fits well with the Biospecimen Age category. |
| companion animals | Biospecimen Organism | HIGH | Companion animals are a type of living biological system |
| molecule subtype | Biospecimen Type | HIGH | A molecule subtype is a specific type of biospecimen, referring to a particular category of molecules. |
| enrollment batch | External_sample_id | MEDIUM | Enrollment batch could refer to a group of patients or samples, but it is not a clear fit for any category. |
| segment | Other | LOW | The term 'segment' is ambiguous and could refer to various concepts in biomedical research, such as a segment of DNA or a segment of a clinical trial. |

#### 3.3.3 Fine-Grained Sub-clustering

Some predefined categories are inherently broad and include heterogeneous label types. For example, the *Biospecimen Type* category contains labels describing tissues, cell types, cell lines, sample preparation procedures, and other biospecimen-related attributes. To support more precise querying and downstream analysis, labels within each category were further grouped into fine-grained subcategories.

Candidate subcategories were generated using the same Llama-3.1-70B-Instruct model with low-temperature decoding (temperature = 0.02). For each main category, the model was given the labels assigned to that category and instructed to produce a concise set of coherent subcategories, with a maximum of eight subcategories per main category. The prompt instructed the model to 1) use short descriptive subcategory names, 2) use concise, descriptive subcategory names representing coherent biological or technical concepts, 3) avoid overly granular one-or two-label groups when broader groupings were appropriate, 4) assign every label to exactly one subcategory and avoid creating a generic ‘Other’ subcategory. For categories with large numbers of labels, the labels were first sub-clustered in batches, and the union of labels from those batch-level results was then re-clustered into a final set of up to eight representative subcategories for the full category.

We then applied a second LLM reassignment prompt for labels omitted from the initial sub-clustering. This follow-up prompt provided the existing subcategory names, representative labels already assigned to each subcategory, and the omitted labels, and instructed the model to assign each omitted label to one of the existing subcategories.

If any labels remained un-clustered after the reassignment step, they were assigned to the most similar existing subcategory using string similarity between the label and the subcategory name or example labels. If an initial LLM response could not be parsed, the script retried the task on smaller label subsets; when necessary, a fallback single-cluster assignment was applied.

After LLM-assisted clustering, we reviewed the 69 LLM-generated subcategories and manually consolidated them into 17 curated subcategories. The consolidation focused on merging biologically similar or overly granular groups while preserving distinctions relevant to dataset discovery and secondary analysis. For example, LLM-generated groups such as *Tissue and Cell Types*, *Cell Sources*, *Cell Features*, and *Cell Subgroups* were merged into broader categories including *Tissue and Cell Types* and *Cell Characteristics*.

Examples of LLM-generated subcategories and consolidated subcategories are listed in the Table 7 and Table 8, and the full list of subcategories is provided in the Supplementary File 6 (“Fine_grained_subcategories_sample_characteristics.xlsx”). This two-tier classification framework, from high-level category assignment to curated subcategory consolidation, ensured complete label coverage, consistent naming, and scalable semantic integration of sample-level metadata across GEO studies.

**Table 7.** Examples of LLM-generated subcategories under the category of Biospecimen Type.

| Subcategory | Label Examples |
| --- | --- |
| Tissue and Cell Types | ['tissue', 'cell type', 'cell line', 'tissue type', 'celltype', 'tissue anatomic site'] |
| Cell and Tissue Sources | ['serum type', 'cell culture', 'cell line source', 'cell type/line', 'primary tissues', 'transplanted organ'] |
| Cell and Tissue Subgroups | ['cell state', 'cell compartment', 'cell subpopulation', 'cell density', 'cell culture', 'primary tissues', 'transplanted organ'] |
| Cell and Tissue Characterization | ['bacteria tissue', 'large cell lung cancer Derived from metastatic site', 'cell subpopulation/marker'] |
| Sample Isolation and Preparation | ['library prep', 'protocol', 'protocol description', 'protocol number', 'study protocol'] |
| Tissue and Cell Features | ['waist_circumference', 'microdissection', 'assayed molecule', 'lymphs', 'line', 'isolation_method'] |
| Small Molecules | ['small molecule'] |

**Table 8.** Examples of consolidated subcategories under the category of Biospecimen Type.

| Consolidated Categories | Label Examples |
| --- | --- |
| Tissue and Cell Types | ['cell stage', 'tissues', 'primary cell line', 'tissue subgroup', 'cell line/tissue'] |
| Tissue and Cell Characteristics | ['cell state', 'cell compartment', 'cell subpopulation', 'cell density', 'primary tissues', 'transplanted organ'] |
| Sample Isolation and Preparation | ['library prep', 'protocol', 'protocol description', 'protocol number', 'cell culture', 'study protocol'] |

## 4. Data Model Development

To semantically capture and represent the retrieved data from GEO, we defined a data model including six primary classes, and associated data properties and object properties. The data model is shown in Figure 2.

**Figure 2.**
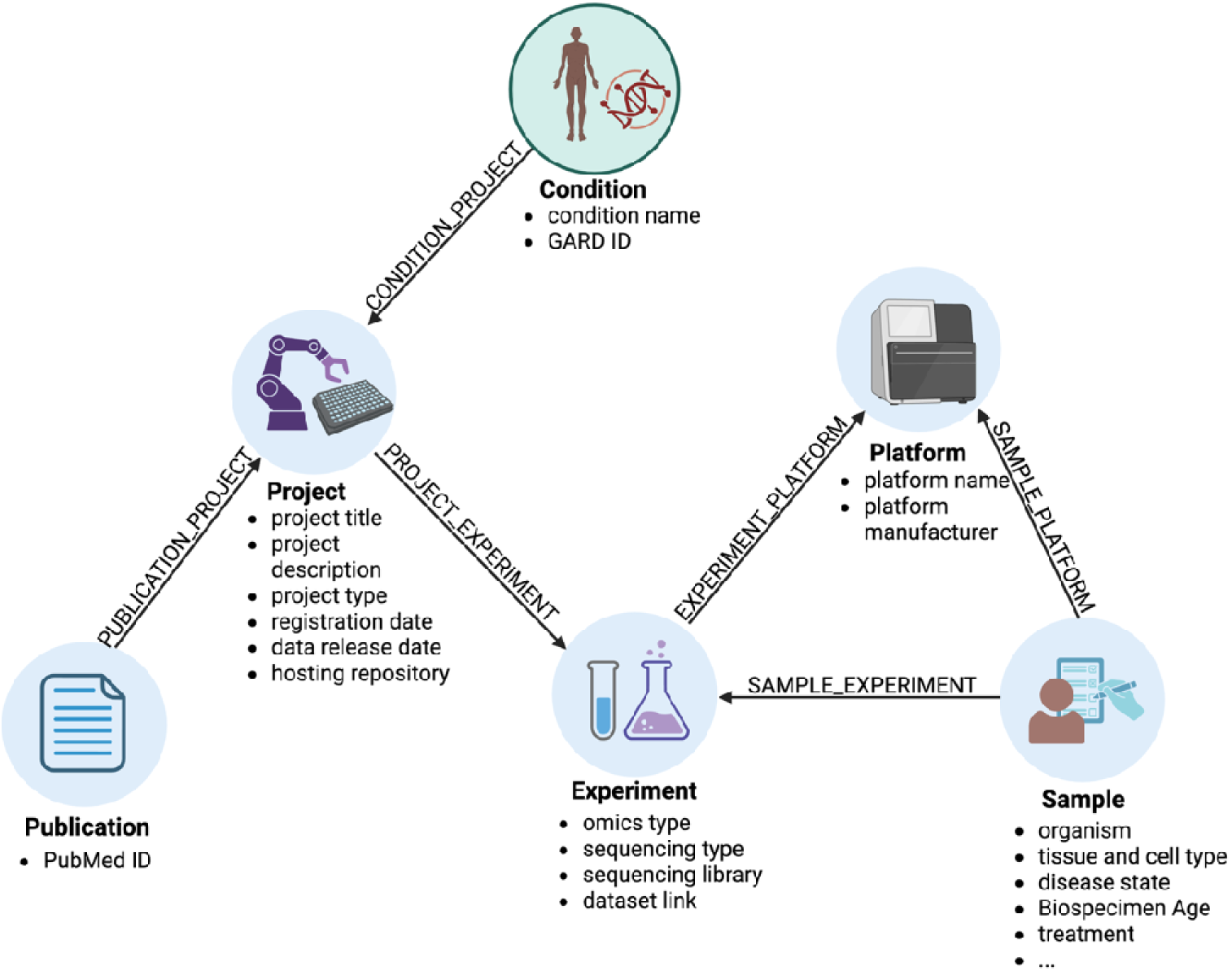
RD-OMICS Data Model. The RD-OMICS knowledge graph consists of six primary classes (Condition, Project, Experiment, Sample, Platform, and Publication) connected through semantic relationships. Representative data properties are shown for each class. Created in BioRender. Sun, S. (2026) https://BioRender.com/yyqsqfu.

### 4.1 Primary Class Definition

We defined six primary classes according to the different types of data we obtained from GEO. *Condition* was defined to capture disease-specific context and serve as the anchor for organizing datasets associated with the same RD. *Project* represents overarching research programs (e.g., BioProjects in GEO). *Experiment* describes distinct omics experiment and the associated data-generate activities. *Sample* represents individual biological specimens that were measured in an *Experiment. Platform* encodes information about the sequencing or microarray platforms used to generate omics data. *Publication* stores bibliographic information (e.g., PubMed ID) describing studies associated with *Projects* and *Experiments*.

### 4.2 Data Property Definition

For each primary class, a list of data properties was created to provide more detailed information about the class. The experiment-and sample-level metadata were captured as the data properties of the *Experiment* and *Sample* classes, respectively. Additional data properties were derived from GEO metadata fields associated with projects, platforms, publications, and disease conditions and attached to corresponding classes, such as, *Project*, *Platform*. Examples are listed in Table 9.

**Table 9.**
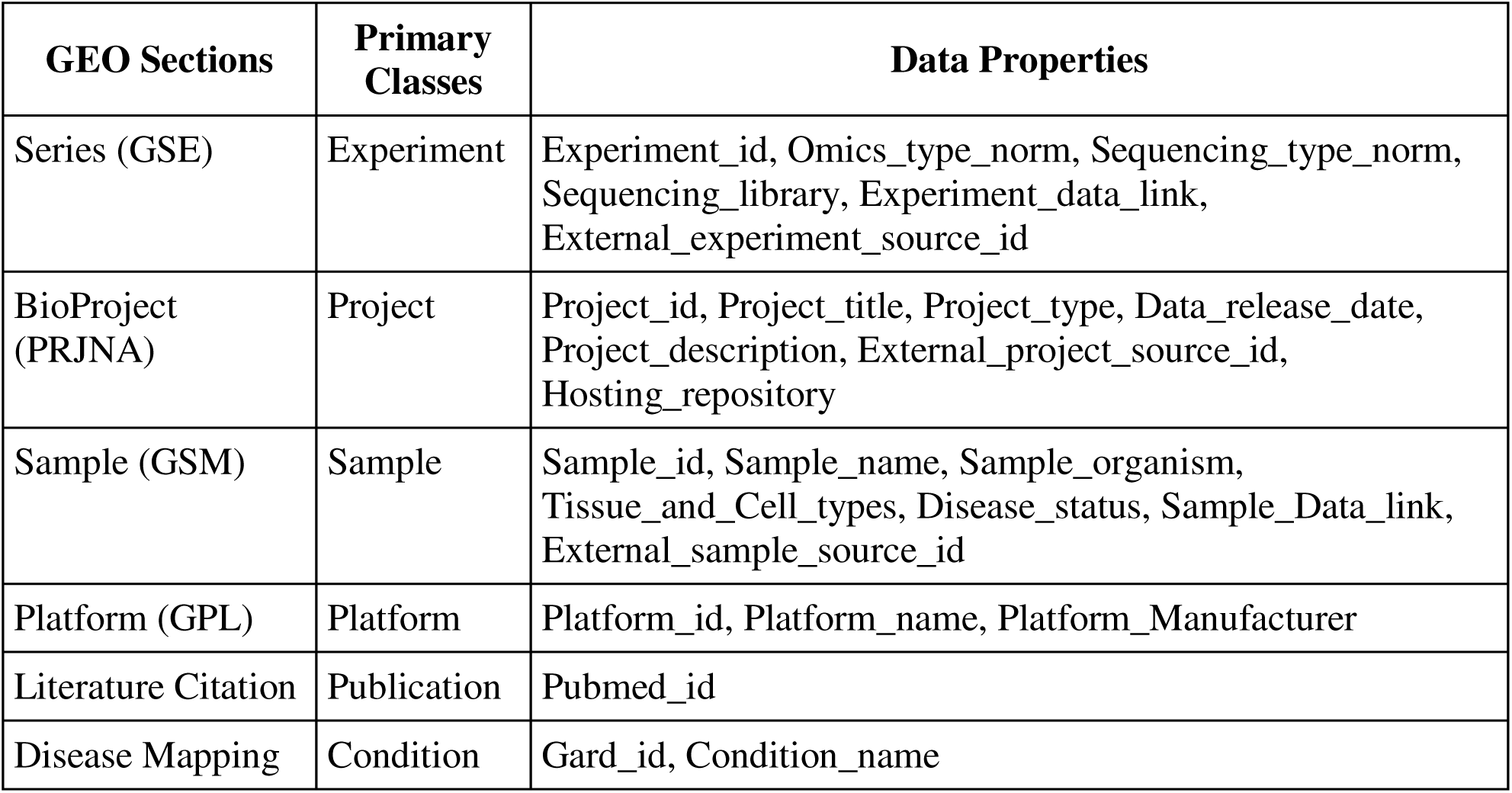
Defined primary classes and associated data properties.

### 4.3 Object Property Definition

We defined object properties to represent semantic relationships among primary classes. The list of object properties is listed in Figure 2. These relationships were designed to organize heterogeneous GEO metadata into a rare disease–centered graph structure that supports efficient dataset discovery, cohort construction, and downstream analyses. For example, SAMPLE_OF links each *Sample* to its corresponding *Experiment*, while PART_OF_PROJECT associates *Experiments* with their parent *BioProject*. Disease-centered relationships, such as ASSOCIATED_WITH_CONDITION, connect projects to standardized rare disease concepts using curated GARD mappings.

## 5. RD-OMICS Development

Based on the data model described above, we populated multi-omics data retrieved from the GEO to a knowledge graph hosted in Memgraph graph database (45). Different types of data have been loaded and represented as the six primary classes as nodes (Figure 2); object properties were applied to establish semantic connections between different nodes as edges, and data properties were attached to corresponding nodes as node properties. The data applied for the knowledge graph construction can be found at https://github.com/ncats/RDAS/tree/master/RDAS_RDOMICS.

## Results

### 1. Results on GEO Data Preparation

Out of the initial list of 194 RDs, a total of 11,049 GSEs were identified for 126 RDs, which is provided in Supplementary File 7 (“RD_GES_list.xlsx”). Among these retrieved GSEs, there are 375,930 individual GSM, 1,578 sequencing and array platforms, 10,938 biological projects.

### 2. Results on Experiment-Level Metadata Normalization

Among the 11,049 experiments, we observed substantial heterogeneity in experiment-level metadata, 316 unique combinations of raw values across the *Data type*, *Library strategy*, and *Extracted molecule* fields. From the rule-based normalization approach described in the Methods, these metadata combinations were mapped to 30 standardized experiment categories. For example, an experiment normalized as Transcriptomics – RNA-seq – ncRNA represents transcriptomics profiling using RNA-seq sequencing on non-coding RNA samples (Table 3). After normalization, the experiments were classified into five high-level omics domains: 8,924 transcriptomics experiments, 1,504 epigenomics experiments, 525 genomics experiments, 76 proteomics experiments, and 20 other omics experiments.

Table 10 summarizes the harmonized results of assay types within each omics domain. RNA-seq was the dominant assay among transcriptomic experiments, accounting for 58.8% of transcriptomics studies, whereas ChIP-seq (42.7%) and ATAC-seq (22.3%) were the most common assay types among epigenomic studies. In contrast, genomic and proteomic experiments showed greater heterogeneity in their assay annotations. Overall, 4,506 experiments (40.8%) were classified as *Other,* primarily because the available GEO metadata were insufficient to determine a more specific sequencing strategy. To fill the gap, we plan to analyze other fields, rather than relying only on *Data type*, *Library strategy*, and *Extracted molecule* in the future study.

**Table 10.** Normalization results on assay types across omics domains.

| **Assay type** | **Epigenomics**<br>**(N=1,504)** | **Genomics**<br>**(N=525)** | **Proteomics**<br>**(N=76)** | **Transcriptomics**<br>**(N=8,924)** | **Other**<br>**(N=20)** | **Overall**<br>**(N=11,049)** |
| --- | --- | --- | --- | --- | --- | --- |
| ATAC-seq | 335 (22.3%) | 0 (0%) | 0 (0%) | 0 (0%) | 0 (0%) | 335 (3.0%) |
| Bisulfite-seq | 91 (6.1%) | 0 (0%) | 0 (0%) | 0 (0%) | 0 (0%) | 91 (0.8%) |
| ChIA-PET | 2 (0.1%) | 0 (0%) | 0 (0%) | 0 (0%) | 0 (0%) | 2 (0.0%) |
| ChIP-seq | 642 (42.7%) | 0 (0%) | 0 (0%) | 0 (0%) | 0 (0%) | 642 (5.8%) |
| DNase-Hypersensitivity | 18 (1.2%) | 0 (0%) | 0 (0%) | 0 (0%) | 0 (0%) | 18 (0.2%) |
| FAIRE-seq | 2 (0.1%) | 0 (0%) | 0 (0%) | 0 (0%) | 0 (0%) | 2 (0.0%) |
| Hi-C assay | 23 (1.5%) | 0 (0%) | 0 (0%) | 0 (0%) | 0 (0%) | 23 (0.2%) |
| MBD-Seq | 3 (0.2%) | 0 (0%) | 0 (0%) | 0 (0%) | 0 (0%) | 3 (0.0%) |
| MeDIP-Seq | 10 (0.7%) | 0 (0%) | 0 (0%) | 0 (0%) | 0 (0%) | 10 (0.1%) |
| MNase-Seq | 3 (0.2%) | 0 (0%) | 0 (0%) | 0 (0%) | 0 (0%) | 3 (0.0%) |
| RNA immune-precipitation | 70 (4.7%) | 0 (0%) | 0 (0%) | 0 (0%) | 0 (0%) | 70 (0.6%) |
| RNA-Seq | 5 (0.3%) | 63 (12.0%) | 0 (0%) | 5,250 (58.8%) | 0 (0%) | 5,318 (48.1%) |
| Metagenomics | 0 (0%) | 1 (0.2%) | 0 (0%) | 0 (0%) | 0 (0%) | 1 (0.0%) |
| Tn-Seq | 0 (0%) | 2 (0.4%) | 0 (0%) | 0 (0%) | 0 (0%) | 2 (0.0%) |
| SELEX | 0 (0%) | 0 (0%) | 1 (1.3%) | 0 (0%) | 0 (0%) | 1 (0.0%) |
| Single cell sequencing | 0 (0%) | 0 (0%) | 0 (0%) | 24 (0.3%) | 0 (0%) | 24 (0.2%) |
| Other | 300 (19.9%) | 459 (87.4%) | 75 (98.7%) | 3650 (40.9%) | 20 (100%) | 4506 (40.8%) |

Table 11 summarizes the distribution of harmonized extracted molecule types across omics domains. As expected, total RNA was the predominant molecule type among transcriptomic experiments (7,792/8,924, 87.3%), whereas genomic DNA was the dominant molecule type in both epigenomic (70.8%) and genomic (72.6%) studies. Protein was almost exclusively associated with proteomic experiments (96.1%). A subset of experiments (n = 665, 6.0%) was classified as multiple molecule, indicating that more than one molecular material and assay type was performed within the same experiment. These results further demonstrate that the harmonized metadata preserved biologically meaningful distinctions in the molecular substrates underlying different omics domains.

**Table 11.** Normalization results on extracted molecule types across omics domains.

| **Extracted molecule** | **Epigenomics**<br>(N=1,504) | **Genomics**<br>(N=525) | **Proteomics**<br>(N=76) | **Transcriptomics**<br>(N=8,924) | **Other**<br>(N=20) | **Overall**<br>(N=11,049) |
| --- | --- | --- | --- | --- | --- | --- |
| total RNA | 61 (4.1%) | 1 (0.2%) | 0 (0%) | 7792 (87.3%) | 0 (0%) | 7854 (71.1%) |
| polyA RNA | 14 (0.9%) | 0 (0%) | 0 (0%) | 930 (10.4%) | 0 (0%) | 944 (8.5%) |
| genomic DNA | 1065 (70.8%) | 381 (72.6%) | 0 (0%) | 0 (0%) | 0 (0%) | 1446 (13.1%) |
| protein | 0 (0%) | 0 (0%) | 73 (96.1%) | 1 (0.0%) | 0 (0%) | 74 (0.7%) |
| nuclear RNA | 0 (0%) | 0 (0%) | 0 (0%) | 21 (0.2%) | 0 (0%) | 21 (0.2%) |
| cytoplasmic RNA | 0 (0%) | 0 (0%) | 0 (0%) | 4 (0.0%) | 0 (0%) | 4 (0.0%) |
| multiple molecule | 361 (24.0%) | 143 (27.2%) | 1 (1.3%) | 160 (1.8%) | 0 (0%) | 665 (6.0%) |
| other | 3 (0.2%) | 0 (0%) | 2 (2.6%) | 16 (0.2%) | 17 (85.0%) | 38 (0.3%) |
| unknown | 0 (0%) | 0 (0%) | 0 (0%) | 0 (0%) | 3 (15.0%) | 3 (0.0%) |

### 3. Results on Sample-Level Metadata Harmonization

Among 375,930 retrieved samples, we identified 5,555 unique submitter-defined labels describing biospecimen and sample characteristics. After multi-phase cleaning and filtering, 2,814 syntactically valid labels were retained for semantic clustering and categorization.

The cleaned labels were first mapped to ten high-level biomedical categories (Table 7) and then further organized into 17 curated subcategories (Table 8). These subcategories captured key dimensions of sample-level metadata, including Biospecimen Type, Sex, Age, Treatment Protocol. This enables unified sample filtering, cohort construction, and visualization in downstream tasks.

Using the hierarchical categorization framework described in the Methods, these labels were first assigned to ten high-level biomedical categories and subsequently consolidated into 17 curated subcategories (Table 12). The resulting subcategories captured key dimensions of sample metadata, including tissue and cell types, disease state, disease progression, treatment information, organism characteristics, sex, race/ethnicity, and treatment regimen. For example, the *Biospecimen Type* category was refined into subcategories, such as *Tissue and Cell Types*, *Cell and Tissue Characteristics*, and *Sample Isolation and Preparation*. Similarly, *Biospecimen Disease Condition* was resolved into more specific subcategories, including *Disease State*, *Disease Characteristics*, *Disease Progression*, *Disease Details*, and *Disease Severity*.

**Table 12.** Categorized Properties and Example Labels of Sample Node.

| Categories | Sub-categories | Label Examples | Total # of Distinct Labels |
| --- | --- | --- | --- |
| Biospecimen Type | Tissue and Cell Types | cell stage, tissues, primary cell line | 149 |
|  | Cell and Tissue Characteristics | serum type, specimen_name, cell culture | 149 |
|  | Sample Isolation and Preparation | library prep, protocol, protocol description | 99 |
| Biospecimen Disease Condition | Disease State | disease state, disease, diagnosis | 136 |
|  | Disease Characteristics | infection group, worm infection, Tumor size | 255 |
|  | Disease Progression | interim pet response, primary/recurrent, pasi_total | 70 |
|  | Disease Details | histological response, hereditary status, Family History | 93 |
|  | Disease Severity | mitotic rate, positive Down screening risk, who class | 3 |
| Biospecimen Age | Biospecimen Age | age, age/gender, time | 85 |
| Treatment | General Treatment Info | treatment, treatment condition, group | 82 |
| Biospecimen Organism | Cell Strain | strain, strain background, strain/background | 43 |
|  | Organism | organism, organism part, cell organism | 33 |
| Biospecimen Sex | Sex | gender, Sex, sex | 24 |
| Biospecimen Race | Race/Ethnic | self_reported_race, race/ethnicity, race_ethnicity | 11 |
|  | Ancestral Origin | ancestry | 1 |
| Treatment Dosage Regimen | Treatment Schedule | dose, immunization dose, vaccination dose | 15 |
|  | Treatment Duration | duration, duration of illness, culture duration | 11 |

This harmonization process transformed heterogeneous GEO submitter-defined metadata into standardized sample-level properties, enabling more consistent sample filtering, cohort construction, visualization and cross-study analysis within RD-OMICS.

## 4. Results on Knowledge Graph Development

To evaluate the content and coverage of RD-OMICS, we performed a graph-level summary analysis of condition-linked experiments, samples, omics modalities, organism annotations, and tissue or cell-type metadata. Sample counts varied substantially across RDs, spanning several orders of magnitude, with most conditions represented by hundreds to thousands of samples and a small number of conditions having much larger sample collections (Figure 3.A). In contrast, most RDs were represented by relatively few experiment records, while only a small number of RDs were associated with hundreds to thousands of experiments (Figure 3.B).

**Figure 3.**
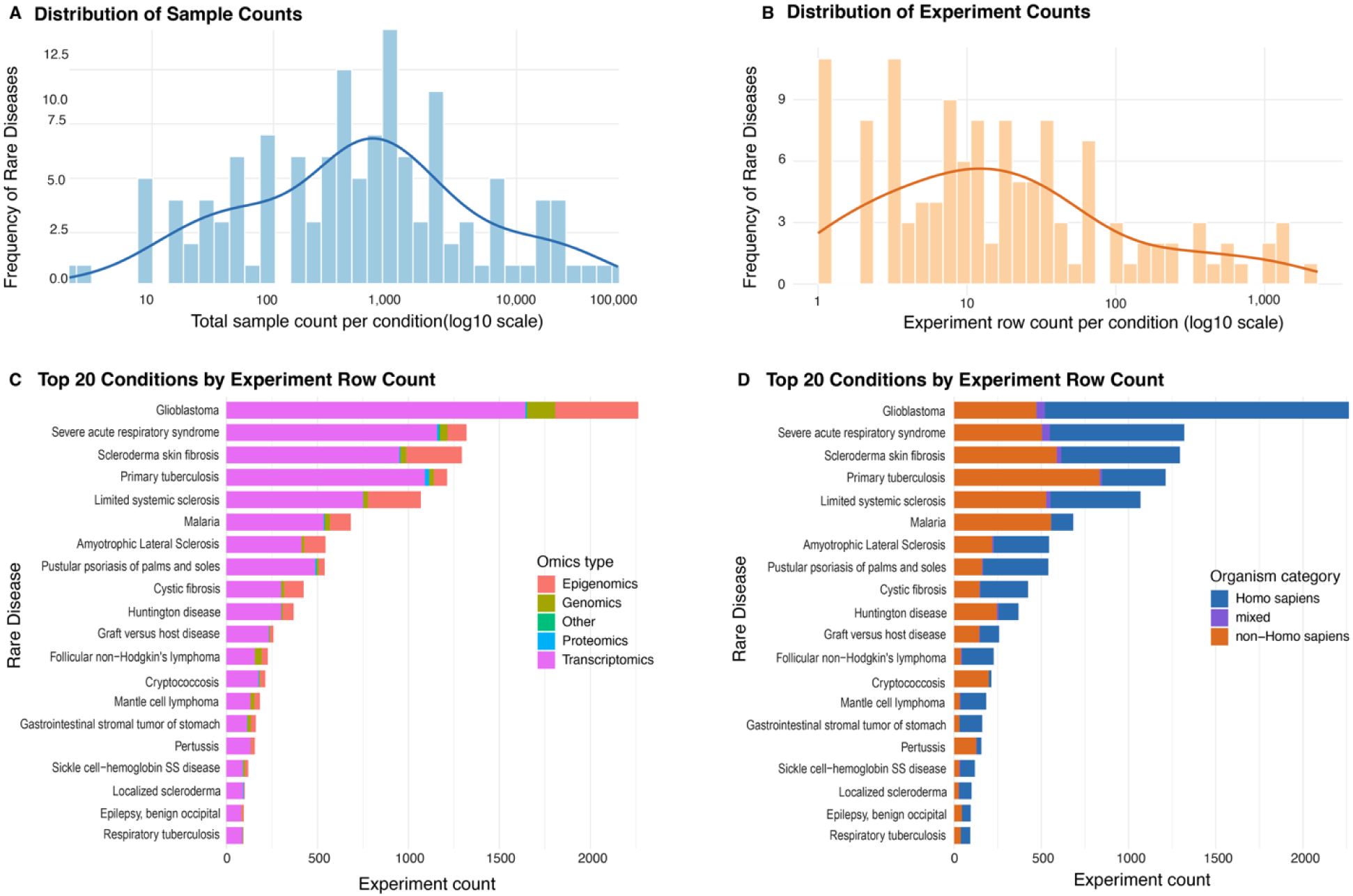
Overview of multi-omics datasets represented in RD-OMICS. (A) Distribution of total sample counts across disease conditions. (B) Distribution of experiment record counts across disease conditions. Histograms are shown with smoothed density curves and log10-scaled count values. (C) Omics modality composition among the top 20 conditions ranked by experiment-condition record count following consolidation of redundant condition labels. (D) Organism composition among the same top 20 conditions. Organism annotations were grouped into *Homo sapiens*, non-*Homo sapiens*, and mixed categories, where mixed indicates studies containing both human and non-human organism annotations.

Among the top 20 conditions by experiment record count, glioblastoma was the most represented condition, followed by severe acute respiratory syndrome, scleroderma skin fibrosis, primary tuberculosis, and limited systemic sclerosis (Figure 3.C). Transcriptomics accounted for the largest share of experiment records across most top conditions, while epigenomics contributed substantially for several highly represented conditions, particularly glioblastoma. Genomics, proteomics, and other omics types were present but represented smaller proportions overall.

Organism-level analysis showed that RD-OMICS includes both human and model-organism studies (Figure 3.D). Among the top 20 conditions, several conditions were dominated by Homo sapiens records (54.4%), including glioblastoma and scleroderma skin fibrosis, whereas others included substantial non-Homo sapiens representation (43.9%), such as primary tuberculosis and severe acute respiratory syndrome. Mixed-species records (1.7%) were present but contributed a smaller fraction. Together, these results demonstrate that RD-OMICS captures a broad range of rare disease-associated omics datasets while also highlighting uneven disease coverage, transcriptomics-heavy representation, and variable use of human versus model-organism data across conditions.

## Case Studies

### 1 RD-OMICS–Enabled Retrieval of GBM Transcriptomic RNA-seq Data

To demonstrate how RD-OMICS facilitates efficient retrieval of omics data for downstream studies, we conducted a GBM-focused case study comparing RD-OMICS based querying with manual dataset discovery. The objective was to identify GEO datasets containing bulk transcriptomic RNA-sequencing data derived from human brain tissue samples from patients with GBM.

### 1.1 Manual Dataset Retrieval

A manual search was performed in GEO using GBM-related keywords and filters to identify relevant human transcriptomics datasets. The search query shown in Query 1, included GBM related terms, restricted results to Homo sapiens, limited records to GEO series, applied the tissue attribute filter, and selected studies annotated as “Expression profiling by high throughput sequencing.” The query returned 826 GEO series (Supplementary File 8, “GEO_GBM_search.pdf”, page 1).

**Query 1**: (“glioblastoma”[MeSH Terms] OR glioblastoma [All Fields]) AND “Homo sapiens”[porgn] AND ((“gse”[Filter] OR “gds”[Filter]) AND “attribute name tissue”[Filter])”) Because GEO metadata are heterogeneous and not uniformly structured, each series required manual inspection to determine:

● Whether the assay was bulk RNA-seq rather than single-cell sequencing
● Whether samples were derived from primary brain tissue rather than cell lines or stem cell models
● Whether the cohort included GBM patients and appropriate controls

Through detailed manual screening, only three GEO series, GSE48865, GSE107559, and GSE149009 met all inclusion criteria.

To determine whether a more restrictive search strategy could reduce the amount of manual review, we performed an additional GEO search using explicit keywords related to bulk RNA-seq, RNA-seq platform technology, and control samples (Query 2).

**Query 2:** ((((((Glioblastoma) OR Glioblastoma) AND Homo sapiens[Organism]) AND bulk RNA-seq) OR RNA-seq[Platform Technology Type]) AND control[Description])

This refined query returned 17 candidate studies (Supplementary File 8, “GEO_GBM_search.pdf”, page 2); however, none satisfied all study inclusion criteria. These results illustrate the difficulty of constructing sufficiently sensitive and specific GEO search queries, as relevant experiment characteristics are distributed across multiple metadata fields and described using heterogeneous submitter-defined terminology.

### 1.2 RD-OMICS–Based Dataset Retrieval

We queried RD-OMICS to retrieve GBM-related omics experiments together with the associated experiment-level metadata, including omics domain, assay type, extracted molecule, as well as sample-level metadata, including tissue and cell type, disease state, organism, sex, age, treatment status.

A total of 2,263 GBM-related omics experiments were retrieved. Using the harmonized metadata fields, we applied structured filters to identify experiments matching the case-study criteria:

● omics domain = transcriptomics
● assay type = RNA-seq
● extracted molecule = total RNA
● tissue and cell type containing the term “tissue”
● exclusion of experiments involving treatment to focus on disease expression profiling studies.

This filtering yielded 15 candidate experiments. Manual verification confirmed that 12 of the 15 experiments satisfied all inclusion criteria. The complete set of GBM-related omics experiments and the filtered results are provided in the Supplementary File 9 (“RD-OMICS_case_study.xlsx”, sheet 1 and 2). Compared with manual GEO searching, using RD-OMICS reduced the number of studies requiring manual inspection from 826 to 15, corresponding to a 98.2% reduction in manual screening burden.

### 1.3 Comparison of Identified Datasets

Using normalized experiment-and sample-level metadata, RD-OMICS filtering retrieved 15 candidate GBM experiments, of which 12 were confirmed by manual review. In comparison, conventional GEO searching required manual inspection of hundreds of GEO series and ultimately identified three eligible datasets. Although the two search strategies were not identical and therefore are not directly comparable, the RD-OMICS workflow substantially reduced the amount of manual review required by enabling structured filtering based on harmonized metadata.

Among the datasets retrieved through RD-OMICS, GSE149009 overlapped with the manually identified results. The remaining two manually identified datasets (GSE48865 and GSE107559) were not retained in the filtered RD-OMICS results, because the tissue and cell type information was either missing from the extracted metadata fields or recorded in fields that are beyond the scope of RD-OMICS.

This case study demonstrates that querying of harmonized experiment-and sample-level metadata in RD-OMICS can substantially improve the efficiency and coverage of dataset discovery, while maintaining high precision. The remaining discrepancies were primarily attributable to incomplete, inconsistent or out-of-scope metadata in the original GEO records, highlighting opportunities for future expansion of the RD-OMICS extraction and harmonization process.

## 2 Multi-Omics Cohort Construction for ALS

To demonstrate the utility of RD-OMICS for cohort construction and multi-omics exploration, we performed an ALS-focused case study. The objective was to retrieve ALS-related omics experiments represented in RD-OMICS and summarize their specimen characteristics and omics distributions using harmonized experiment-and sample-level metadata.

### 2.1 Cohort Retrieval

We searched experiments related to ALS in RD-OMICS. For each experiment, we extracted harmonized experiment-level and sample-level metadata, including omics domain, assay type, tissue and cell type, disease state, organism, and platform. These metadata fields were then used to characterize the available ALS-related datasets by omics modality and biospecimen context. The resulting query returned 544 ALS-related omics experiments (Supplementary File 9, “RD-OMICS_case_study.xlsx”, sheet 3). Representative examples are shown in Table 13 to illustrate how RD-OMICS supports ALS cohort construction by linking disease condition, omics modality, sequencing assay, platform, organism, and biospecimen information within a single searchable resource.

**Table 13.**
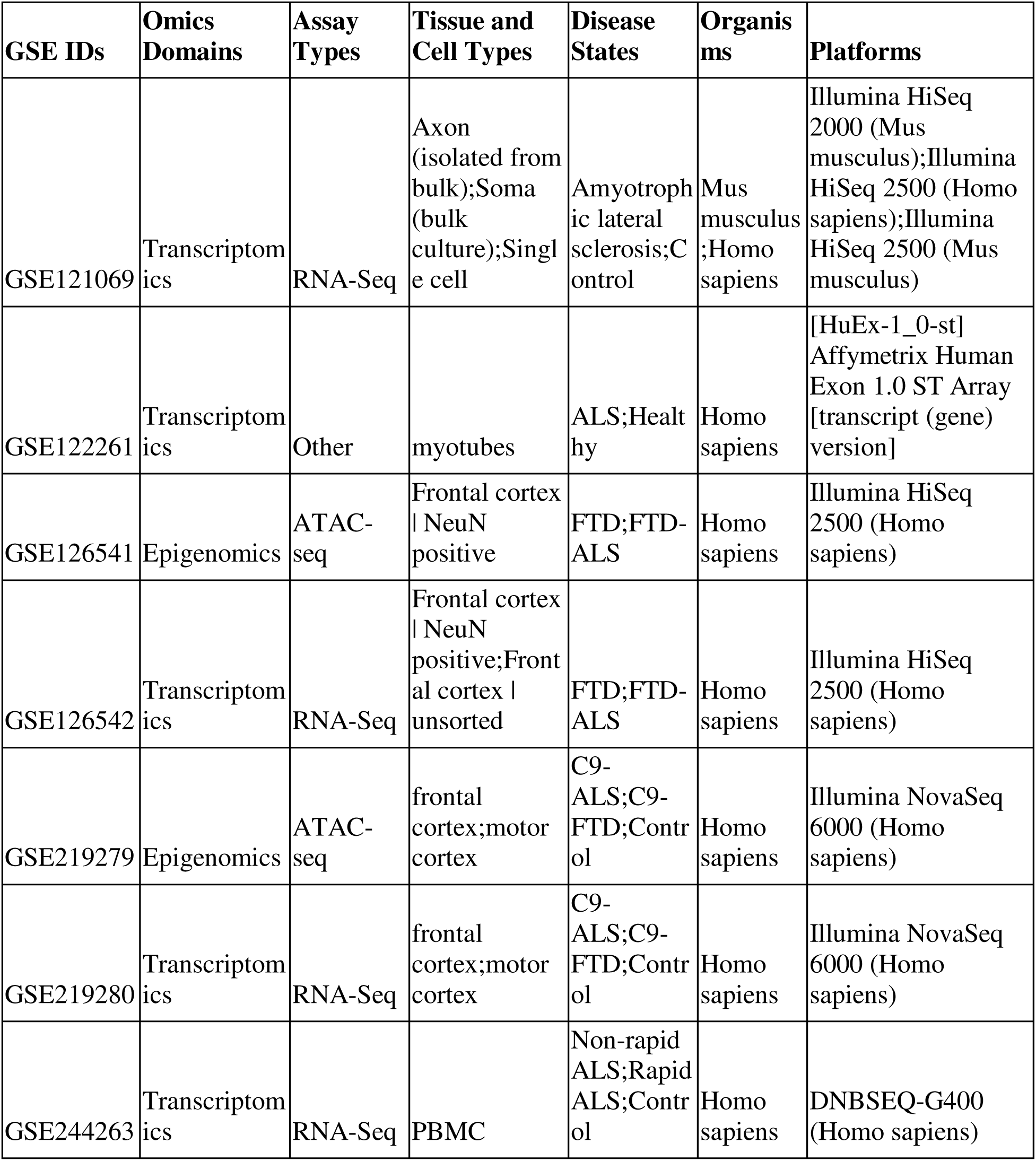
ALS-related omics data retrieved from RD-OMICS.

### 2.2 Specimen Distribution

Applying harmonized sample-level metadata, we summarized the distribution of tissue and cell types across the ALS cohort. Figure 4.A presents the 20 most frequently represented specimen categories. Motor neurons and spinal cord tissue were among the most common studied specimen types, consistent with the central role of moto neuron degeneration and spinal cord pathology in ALS. Other frequently represented categories included induced pluripotent stem cell (iPSC)-derived models, cortical tissues, stem cells, and peripheral blood mononuclear cells (PBMCs), reflecting the diversity of experimental systems used to study ALS biology across both disease-relevant neural tissues and accessible peripheral biospecimens.

**Figure 4.**
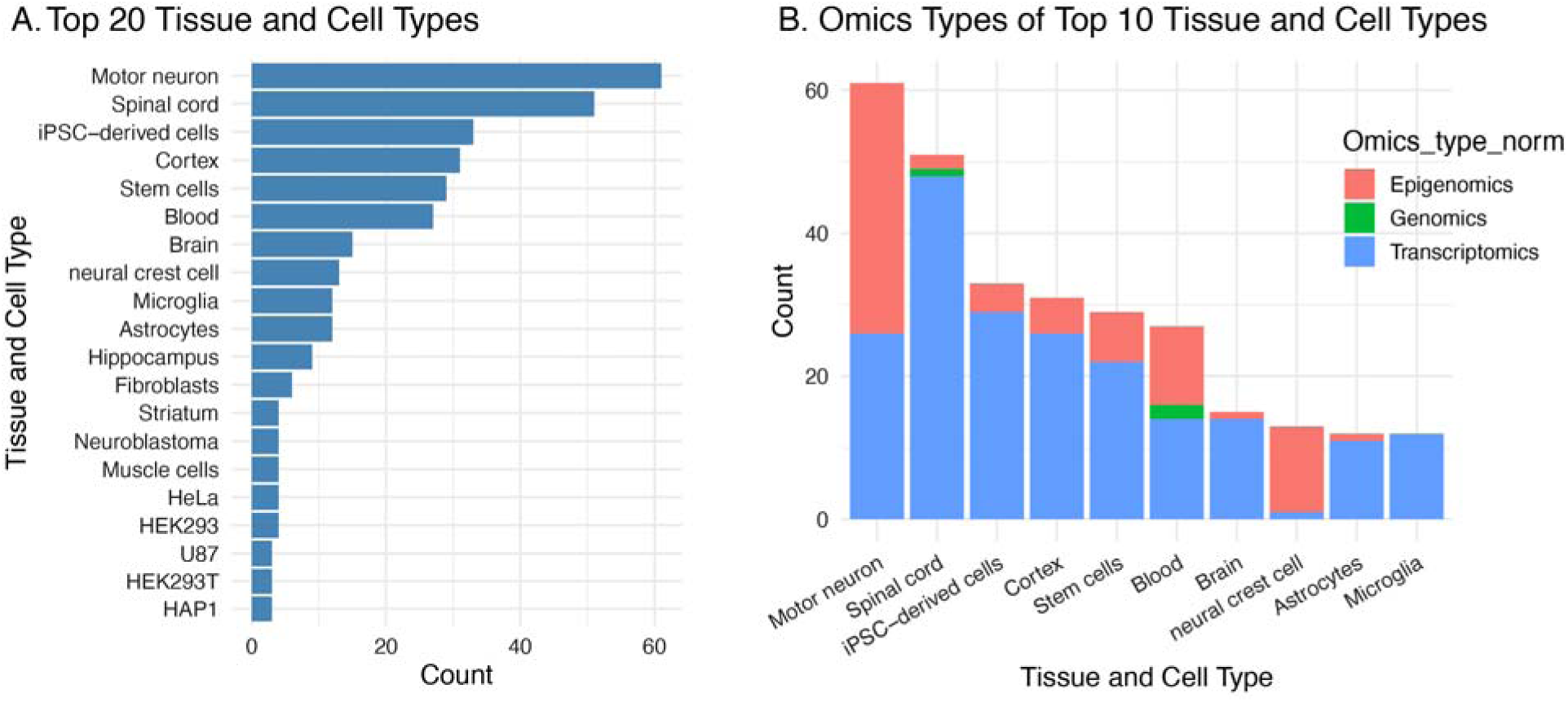
Distribution of specimen types and omics modalities in the ALS cohort. (A) Bar chart showing the 20 most frequently represented tissue and cell types among ALS-related omics experiments in RD-OMICS. Counts indicate the number of experiments associated with each specimen category based on harmonized sample-level metadata. (B) Stacked bar chart showing the distribution of normalized omics modalities across the ten most frequently represented tissue and cell types. Transcriptomics was the predominant modality across most specimen categories, whereas epigenomics and genomics were more selectively represented.

### 2.3 Omics Modality Distribution

We then examined omics modality distribution across the most frequently represented ALS specimen types (Figure 4.B). Transcriptomics was the most dominant modality and was broadly represented across diverse specimen categories, supporting its central role in ALS molecular profiling. Epigenomic studies such as ATAC-seq, were also present but were concentrated in a smaller number of specimen types, particularly motor neurons and neural crest cell–related models. This distribution highlights both the breadth of transcriptomic data available for ALS and the more targeted use of epigenomic approaches in selected disease-relevant cellular systems.

Together, this case study illustrates that RD-OMICS enables efficient construction of disease-specific multi-omics cohorts and systematic summarization of experiment-and sample-level attributes without manual inspection of individual GEO records.

## 3 Drug Repurposing Application in ALS

To further demonstrate the downstream analytical utility of RD-OMICS, we conducted an ALS-focused drug repurposing case study using the data identified from RD-OMICS. Specifically, we applied our previous pipeline (16) to ALS omics data retrieved from RD-OMICS to support therapeutic discovery.

### 3.1 Dataset Selection

From the ALS cohort retrieved in above Case Study 2, we applied the following inclusion criteria:

- RNA-seq data
- Human ALS samples with matched healthy controls
- Adequate sample size for differential expression analysis

We identified one GEO series (GSE234297) (46) that met these criteria. The structured metadata enabled direct identification of this dataset without manual review.

### 3.2 Differential Expression and Signature Matching

Using the data access link as one data property stored in the *Experiment* node, we downloaded the corresponding ALS RNA-seq dataset (raw gene count and offset matrix; Illumina HiSeq 2000 platform) from GSE234297. Differential expression analysis was performed to derive an ALS disease gene expression signature, with differentially expressed genes defined as those with FDR < 0.05 and fold change > 1.5 or < 0.67. This analysis identified 245 differentially expressed (DE) genes, including 85 upregulated and 160 downregulated genes (Figure 5.A).

**Figure 5.**
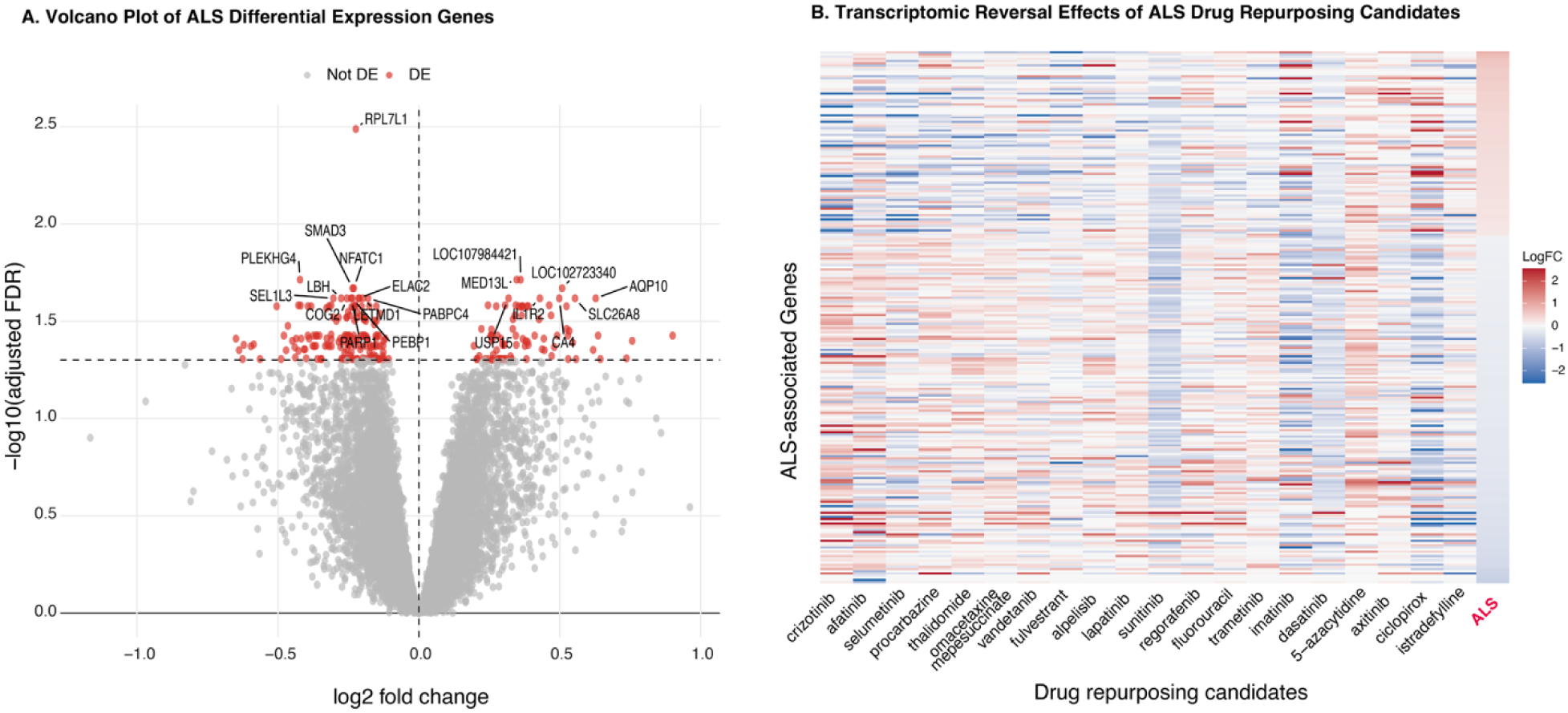
Transcriptomic reversal patterns of candidate drugs for ALS. (A) Volcano plot showing differential gene expression analysis of ALS vs. healthy control samples from RNA-seq dataset GSE234297. Red points indicate significantly differentially expressed genes. (B) Heatmap showing transcriptomic reversal patterns of the top-ranked candidate drugs relative to the ALS disease signature. Rows represent ALS-associated differentially expressed genes, and columns represent drug-induced transcriptional signatures from iLINCS perturbation experiments. Color intensity indicates the direction and magnitude of gene expression changes (red: upregulation; blue: downregulation). Drug signatures exhibiting expression patterns opposite to the ALS disease signature were prioritized as candidate repurposing compounds.

We then queried four iLINCS drug-perturbation signature libraries (47) to identify compounds whose induced gene expression profiles were inversely correlated with the ALS disease signature, suggesting their potential to reverse ALS-associated gene expression changes (Figure 5.B). Across the four libraries, 2,069 drug-induced gene expression signatures exhibited negative correlations with the ALS profile. These signatures corresponded to 731 unique compounds, including 210 FDA-approved drugs (Supplementary File 9, “RD-OMICS_case_study.xlsx”, sheet 4). A future study will be performed to identify the promising drug candidates for ALS by prioritizing and evaluating the 210 drugs.

The case studies demonstrate that RD-OMICS enables rapid identification of appropriate omics datasets and standardized retrieval of experiment-and sample-level metadata. In turn, RD-OMICS reduces manual curation burden and facilitates reproducible, disease-specific computational analyses.

## Discussion

RD research increasingly depends on integrative analysis of omics datasets yet the process of identifying datasets for re-analysis or meta-analyses takes extensive manual curation, largely due to inconsistencies in meta-data reporting. To address this, RD-OMICS integrates RD–related omics datasets into a harmonized knowledge graph supported through systematic metadata harmonization, normalization and categorization. By transforming heterogeneous GEO metadata into harmonized and structured experiment-and sample-level properties, RD-OMICS provides a consistent representation of multi-omics studies across rare diseases, and enables systematic exploration across diseases, omics modalities, organisms, and biospecimen contexts.

## 1. Advantages for RD-OMICS

Several public platforms provide access to omics datasets relevant to rare diseases, including GEO (19) and ArrayExpress (26), the Genome Sequence Archive (48), as well as disease-focused resources and omics platforms such as DiSignAtlas (49). DiSignAtlas, for example, aggregates transcriptomic datasets to generate standardized disease signatures to support comparative analysis across conditions. Besides, large consortium resources, including the NIH Common Fund Data Ecosystem (50), Clinical Proteomic Tumor Analysis Consortium (51), and the *All of Us* Research Program (30), also provide valuable omics datasets through program-specific data portals. However, many existing resources retain heterogeneous metadata structures, making it difficult to identify relevant experiments without extensive manual review. RD-OMICS complements these resources by harmonizing experiment-and sample-level metadata and representing them within a graph-based data model. This structure enables attribute-based querying across studies using biologically meaningful properties, such as disease condition, omics modality, sequencing assay, organism, tissue or cell type, and disease state. In contrast to the manual and often unstructured exploration of individual repository records, RD-OMICS provides a semantic, integrated inventory of RD omics datasets. It enhances the discoverability, interoperability, and analytical usability of publicly available omics data for rare disease research. The utility of RD-OMICS was evaluated through several case studies to illustrate it provides a structured, query-ready resource that can accelerate rare disease omics dataset discovery, cohort construction, and secondary computational analysis.

## 2 Limitation of RD-OMICS

Despite these advances, we acknowledge several limitations. First, identifying GEO datasets for rare diseases remains challenging. This is largely due to inconsistent disease naming across GEO records, incomplete submitter-provided disease annotations in which disease context appears only in free-text descriptions rather than structured data. Future improvements will require more sophisticated disease entity recognition and ontology mapping approaches that can integrate multiple biomedical controlled vocabularies and contextual information from study titles, summaries, protocols, and associated publications.

Second, although many assay related metadata have been successfully standardized, some assay distinctions remain difficult to infer from GEO metadata alone. In particular, differentiating bulk RNA-seq from single-cell RNA-seq can be challenging when records are broadly annotated as RNA-seq but lack explicit protocol or library-preparation information. In such cases, accurate classification may require review of detailed GEO descriptions or associated publications. This limitation reflects the incomplete and inconsistent nature of repository metadata and highlights the need to incorporate additional text-mining and publication-derived evidence in future versions of RD-OMICS.

Third, heterogeneity in sample-level metadata continues to limit fully automated categorization. Although LLM-assisted classification improved coverage and semantic interpretation of GSM *Characteristics* labels, important properties, including tissue and cell type or disease state, were still missing, inconsistently labeled, or recorded outside the extracted fields in some GEO submissions. For example, in the GBM case study, two manually identified datasets were not recovered through RD-OMICS filtering because tissue or cell type information was absent from the structured *Characteristics* fields or embedded in unexpected fields (e.g., sample identifiers). Similarly, the ALS cohort analysis required additional manual review to reconcile inconsistent submitter-provided tissue and cell type annotations. These examples underscore that the performance of automated harmonization depends strongly on the completeness and quality of the source metadata.

## 4 Future Work

Future development of RD-OMICS will focus on addressing current limitations and expanding its scope, depth and utility for RD research. One major priority is to incorporate richer contextual information into metadata harmonization and categorization. In particular, more systematic LLM-based approaches could leverage additional text sources, including experiment descriptions, sample processing details, library preparation protocols, and associated publications, to improve metadata interpretation, disease entity alignment, and assay classification. Incorporating this contextual reasoning would enable RD-OMICS to generate more accurate experiment-level metadata and better classify complex experimental designs, such as distinguishing bulk from single-cell sequencing studies.

A second important direction is to expand the range of omics data sources beyond GEO. While GEO provides a valuable foundation for transcriptomics and other functional genomics datasets, RD omics data are distributed across multiple public repositories and institutional resources. Future versions of RD-OMICS will aim to integrate additional data sources such as the Sequence Read Archive, dbGaP, BioProject, ArrayExpress, PRIDE, MetaboLights, and other disease-or domain-specific repositories. Broadening the data source coverage will increase dataset completeness, reduce repository-specific bias, and enable more comprehensive discovery of RD-relevant omics studies across genomics, transcriptomics, epigenomics, proteomics, metabolomics, and single-cell modalities.

Another key priority is to broaden disease coverage beyond the current set of 194 selected RDs. Future development will seek to include a larger and more diverse set of rare diseases by aligning disease mentions to standardized vocabularies and ontologies, such as MONDO, Orphanet, and OMIM. This expansion will improve representation of under-studied and underrepresented rare diseases, support cross-disease comparisons, and enable researchers to identify shared molecular mechanisms across phenotypically or genetically related conditions. Expanding disease coverage will also make RD-OMICS more useful for rare disease communities where data are sparse, fragmented, or difficult to discover.

Future versions of RD-OMICS will also aim to generate standardized experiment-level summaries that provide concise overviews of dataset characteristics, including disease context, biospecimen source, organism, omics modality, sequencing assay, and key experimental design features. Such summaries would enable prompt assessment of study relevance and improve dataset discovery for downstream rare disease research.

Another important direction is to extend harmonization beyond meta-data to the data level. RD-OMICS currently focuses on harmonizing experimental-and sample-level metadata; however, meta-analyses of omics datasets require harmonization of the underlying molecular measurements. This is substantially more challenging because omics data types differ in measurement scale, experimental design, preprocessing requirements, and analytical pipelines. Integrating genomic variants, transcript abundance profiles, epigenomic signals, proteomic measurements, and other molecular data will require cross-omics harmonization and standardization strategies and scalable computational approaches. Although challenging, data-level harmonization will be essential for enabling direct comparative analyses across rare diseases and supporting advanced machine learning and AI-driven discovery applications.

In addition, future RD-OMICS development could incorporate quality assessment and provenance tracking to improve data reliability and reuse. Standardized quality indicators, such as sample size, case-control balance, biospecimen type, assay platform, availability of raw data, and completeness of metadata, would help users assess whether a dataset is suitable for specific analytical purposes. Capturing provenance information, including original repository identifiers, publication links, processing history, and ontology mapping decisions, would further enhance transparency, reproducibility, and trustworthiness.

## 7. Conclusion

RD-OMICS provides a scalable and semantically structured resource for organizing heterogeneous RD omics datasets into a unified knowledge graph. By systematically harmonizing experiment-and sample-level metadata, RD-OMICS transforms fragmented repository records into queryable, interoperable data assets, thereby facilitating secondary analysis or meta-analysis efforts. Overall, RD-OMICS thus accelerates omics cohort building for biomarker discovery, mechanism investigation, therapeutic hypothesis generation, and data-driven translational research for rare diseases.

## Data availability

Data and scripts developed in this manuscript are available at https://github.com/ncats/RDAS/tree/master/RDAS_RDOMICS.

## Supporting information

RD_List.xlsx

Experiment_metadata_normalization_rules.xlsx

Term_definition_source.docx

Category_reference_information.csv

Semantic_categorization_sample_characteristics.xlsx

Fine_grained_subcategories_sample_characteristics.xlsx

RD_GES_list.xlsx

GEO_GBM_search.pdf

RD-OMICS_case_study.xlsx

## Acknowledgement

This research was supported by the Intramural Research Program (ZIA TR000548 and ZICTR000410) of the National Institutes of Health (NIH) and the High Value Dataset Program from Office of Data Science Strategy (ODSS)/NIH. The contributions of the NIH author(s) were made as part of their official duties as NIH federal employees, are in compliance with agency policy requirements, and are considered Works of the United States Government. However, the findings and conclusions presented in this paper are those of the author(s) and do not necessarily reflect the views of the NIH or the U.S. Department of Health and Human Services.

## Contributions

HW: led the development and implementation of RD-OMICS and wrote the manuscript; SS: led metadata harmonization, performed case studies and wrote the manuscript; EM: provided technical guidance as well as comments/revisions to the manuscript; QZ: conceived and supervised this study and wrote the manuscript. All authors reviewed and approved the manuscript.

