## Supplementary material for "RD-OMICS: An Integrative Multi-Omics Data Inventory in Rare Diseases": Term_definition_source.docx

| **Term** | **Source Vocabulary** | **Source link** | **Definition** |
| --- | --- | --- | --- |
| Omics (type) | EDAM | [link](https://bioportal.bioontology.org/ontologies/EDAM?p=classes&conceptid=http%3A%2F%2Fedamontology.org%2Ftopic_3391) | The collective characterisation and quantification of pools of biological molecules that translate into the structure, function, and dynamics of an organism or organisms. |
| Genomics | EDAM | [link](https://bioportal.bioontology.org/ontologies/EDAM?p=classes&conceptid=http%3A%2F%2Fedamontology.org%2Ftopic_3391) | Whole genomes of one or more organisms, or genomes in general, such as meta-information on genomes, genome projects, gene names etc. |
| Transcriptomics | EDAM | [link](https://bioportal.bioontology.org/ontologies/EDAM?p=classes&conceptid=http%3A%2F%2Fedamontology.org%2Ftopic_3391) | The analysis of transcriptomes, or a set of all the RNA molecules in a specific cell, tissue etc. |
| Proteomics | EDAM | [link](https://bioportal.bioontology.org/ontologies/EDAM?p=classes&conceptid=http%3A%2F%2Fedamontology.org%2Ftopic_3391) | Proteomics includes any methods (especially high-throughput) that separate, characterize and identify expressed proteins such as mass spectrometry, two-dimensional gel electrophoresis and protein microarrays, as well as in-silico methods that perform proteolytic or mass calculations on a protein sequence and other analyses of protein production data, for example in different cells or tissues. Includes metaproteomics: proteomics analysis of an environmental sample. Protein and peptide identification, especially in the study of whole proteomes of organisms. |
| Metabolomics | EDAM | [link](https://bioportal.bioontology.org/ontologies/EDAM?p=classes&conceptid=http%3A%2F%2Fedamontology.org%2Ftopic_3391) | The systematic study of metabolites, the chemical processes they are involved, and the chemical fingerprints of specific cellular processes in a whole cell, tissue, organ or organism. |
| Lipidomics | EDAM | [link](https://bioportal.bioontology.org/ontologies/EDAM?p=classes&conceptid=http%3A%2F%2Fedamontology.org%2Ftopic_3391) | Lipids and their structures. |
| Sequencing (type) | EDAM;  ORNASEQ | [link](https://bioportal.bioontology.org/ontologies/EDAM/?p=classes&conceptid=http%3A%2F%2Fedamontology.org%2Ftopic_3168&lang=en#visualization)  [link2](https://bioportal.bioontology.org/ontologies/ORNASEQ?p=classes&conceptid=http%3A%2F%2Fpurl.obolibrary.org%2Fobo%2FOBI_0600047) | 1. The determination of complete (typically nucleotide) sequences, including those of genomes (full genome sequencing, de novo sequencing and resequencing), amplicons and transcriptomes. 2. Sequencing assay: the use of a chemical or biochemical means to infer the sequence of a biomaterial |
| Assay (type) | ORNASEQ | [link](https://bioportal.bioontology.org/ontologies/ORNASEQ?p=classes&conceptid=http%3A%2F%2Fpurl.obolibrary.org%2Fobo%2FOBI_0000070) | Assay: A planned process with the objective to produce information about the material entity that is the evaluant, by physically examining it or its proxies. |
| microarray | NCIT | [link](https://bioportal.bioontology.org/ontologies/NCIT/?p=classes&lang=en&conceptid=http%3A%2F%2Fncicb.nci.nih.gov%2Fxml%2Fowl%2FEVS%2FThesaurus.owl%23C44282&jump_to_nav=true) | A piece of glass or plastic on which different samples have been affixed at separate locations in an ordered manner thus forming a microscopic array. The samples are usually DNA fragments but may also be antibodies, other proteins, or tissues. |
| RNA-seq | EDAM | [link](https://bioportal.bioontology.org/ontologies/EDAM/?p=classes&conceptid=http%3A%2F%2Fedamontology.org%2Ftopic_3168&lang=en#visualization) | This includes small RNA profiling (small RNA-Seq), for example to find novel small RNAs, characterize mutations and analyze expression of small RNAs. A topic concerning high-throughput sequencing of cDNA to measure the RNA content (transcriptome) of a sample, for example, to investigate how different alleles of a gene are expressed, detect post-transcriptional mutations or identify gene fusions. |
| LC-MS/MS | NCIT | [link](https://bioportal.bioontology.org/ontologies/NCIT?p=classes&conceptid=http%3A%2F%2Fncicb.nci.nih.gov%2Fxml%2Fowl%2FEVS%2FThesaurus.owl%23C122168) | Liquid Chromatography/Tandem Mass Spectrometry: An analytical technique wherein liquid chromatography is coupled to tandem mass spectrometry in order to separate, identify, and quantify substances in a sample. |
| NGS | EDAM | [link](https://bioportal.bioontology.org/ontologies/EDAM/?p=classes&lang=en&conceptid=http%3A%2F%2Fedamontology.org%2Fdata_3165&jump_to_nav=true) | Next generation sequencing, NGS experiment: sequencing experiment, including samples, sampling, preparation, sequencing, and analysis. it is a Synonym to “Sequencing”. |
| Sequencing library | NCIT;  ORNASEQ | [link](https://bioportal.bioontology.org/ontologies/NCIT?p=classes&conceptid=http%3A%2F%2Fncicb.nci.nih.gov%2Fxml%2Fowl%2FEVS%2FThesaurus.owl%23C148073)  [link2](https://bioportal.bioontology.org/ontologies/ORNASEQ/?p=classes&conceptid=http%3A%2F%2Fpurl.obolibrary.org%2Fobo%2FOBI_0000711) | 1. A collection of double stranded DNA fragments flanked by oligonucleotide sequence adapters to enable their analysis by high-throughput sequencing. 2. is a process which results in the creation of a library from fragments of DNA using cloning vectors or oligonucleotides with the role of adaptors. |
| whole genome sequencing | EDAM | [link](https://bioportal.bioontology.org/ontologies/EDAM/?p=classes&conceptid=http%3A%2F%2Fedamontology.org%2Ftopic_3168&lang=en#visualization) | Laboratory technique to sequence the complete DNA sequence of an organism's genome at a single time. |
| metagenomic sequencing | EDAM | [link](https://bioportal.bioontology.org/ontologies/EDAM/?p=classes&conceptid=http%3A%2F%2Fedamontology.org%2Ftopic_3168&lang=en#visualization) | Approach which samples, in parallel, all genes in all organisms present in a given sample, e.g. to provide insight into biodiversity and function. |
| exome sequencing | EDAM | [link](https://bioportal.bioontology.org/ontologies/EDAM/?p=classes&conceptid=http%3A%2F%2Fedamontology.org%2Ftopic_3168&lang=en#visualization) | Exome sequencing is considered a cheap alternative to whole genome sequencing. Laboratory technique to sequence all the protein-coding regions in a genome, i.e., the exome. |
| total RNA | NCIT | [link](https://bioportal.bioontology.org/ontologies/NCIT?p=classes&conceptid=http%3A%2F%2Fncicb.nci.nih.gov%2Fxml%2Fowl%2FEVS%2FThesaurus.owl%23C163995) | A biological sample comprised of all of the RNA collected from an experimental subject. |
| mRNA | SO | [link](https://bioportal.bioontology.org/ontologies/SO/?p=classes&conceptid=http%3A%2F%2Fpurl.obolibrary.org%2Fobo%2FSO_0000233&lang=en#details) | Messenger RNA is the intermediate molecule between DNA and protein. It includes UTR and coding sequences. It does not contain introns. An mRNA does not contain introns as it is a processed_transcript. The equivalent kind of primary_transcript is protein_coding_primary_transcript (SO:0000120) which may contain introns. This term is mapped to MGED. Do not obsolete without consulting MGED ontology. |
| ncRNA | SO | [link](https://bioportal.bioontology.org/ontologies/SO/?p=classes&conceptid=http%3A%2F%2Fpurl.obolibrary.org%2Fobo%2FSO_0000233&lang=en#details) | (noncoding RNA) An RNA transcript that does not encode for a protein rather the RNA molecule is the gene product. A ncRNA is a processed_transcript, so it may not contain parts such as transcribed_spacer_regions that are removed in the act of processing. |
| single cell sequencing | EDAM | [link](https://bioportal.bioontology.org/ontologies/EDAM/?p=classes&conceptid=http%3A%2F%2Fedamontology.org%2Ftopic_3168&lang=en#visualization) | Combined with NGS (Next Generation Sequencing) technologies, single-cell sequencing allows the study of genetic information (DNA, RNA, epigenome...) at a single cell level. It is often used for differential analysis and gene expression profiling. |
| Platform | GEO | [link](https://www.ncbi.nlm.nih.gov/geo/info/platform.html) | Platforms in GEO refer to the technology or array used to generate the gene expression data. A platform represents the set of probes or features on an array that capture gene expression levels. |
| Biospecimen | NCIT | [link](https://bioportal.bioontology.org/ontologies/NCIT/?p=classes&lang=en&conceptid=http%3A%2F%2Fncicb.nci.nih.gov%2Fxml%2Fowl%2FEVS%2FThesaurus.owl%23C70699&jump_to_nav=true) | Any material sample taken from a biological entity for testing, diagnostic, propagation, treatment or research purposes, including a sample obtained from a living organism or taken from the biological object after halting of all its life functions. Biospecimen can contain one or more components including but not limited to cellular molecules, cells, tissues, organs, body fluids, embryos, and body excretory products. |
| Biospecimen Type | NCIT | [link](https://bioportal.bioontology.org/ontologies/NCIT?p=classes&conceptid=http%3A%2F%2Fncicb.nci.nih.gov%2Fxml%2Fowl%2FEVS%2FThesaurus.owl%23C70713) | The type of a material sample taken from a biological entity for testing, diagnostic, propagation, treatment or research purposes. This includes particular types of cellular molecules, cells, tissues, organs, body fluids, embryos, and body excretory substances. |
| Biospecimen Disease Condition | NCIT, DOID | [Link1](https://bioportal.bioontology.org/ontologies/MONDO/?p=classes&lang=en&conceptid=http%3A%2F%2Fpurl.obolibrary.org%2Fobo%2FMONDO_0000001&jump_to_nav=true);  [Link2](https://bioportal.bioontology.org/ontologies/DOID/?p=classes&lang=en&conceptid=http%3A%2F%2Fpurl.obolibrary.org%2Fobo%2FDOID_4&jump_to_nav=true); | A disease is a disposition to undergo pathological processes that exists in an organism because of one or more disorders in that organism. |
| Biospecimen Age | NCIT; OBI | [Link1](https://bioportal.bioontology.org/ontologies/NCIT/?p=classes&lang=en&conceptid=http%3A%2F%2Fncicb.nci.nih.gov%2Fxml%2Fowl%2FEVS%2FThesaurus.owl%23C25150&jump_to_nav=true);  [Link2](https://bioportal.bioontology.org/ontologies/OBI/?p=classes&lang=en&conceptid=http%3A%2F%2Fpurl.obolibrary.org%2Fobo%2FPATO_0000011&jump_to_nav=true); | How long something has existed; elapsed time since birth. ;  A time quality inhering in a bearer by virtue of how long the bearer has existed. |
| Treatment | OBI | [link](https://bioportal.bioontology.org/ontologies/OBI/?p=classes&lang=en&conceptid=http%3A%2F%2Fpurl.obolibrary.org%2Fobo%2FOGMS_0000090&jump_to_nav=true) | A planned process whose completion is hypothesized by a health care provider to eliminate, prevent, or alleviate a disorder, the signs and symptoms of a disorder, or a pathological process |
| Biospecimen Organism | BCO | [link](https://bioportal.bioontology.org/ontologies/BCO/?p=classes&lang=en&conceptid=http%3A%2F%2Frs.tdwg.org%2Fdwc%2Fterms%2FOrganism&jump_to_nav=true) | A particular organism or defined group of organisms considered to be taxonomically homogeneous. |
| Biospecimen Sex | NCIT | [link](https://bioportal.bioontology.org/ontologies/NCIT/?p=classes&lang=en&conceptid=http%3A%2F%2Fncicb.nci.nih.gov%2Fxml%2Fowl%2FEVS%2FThesaurus.owl%23C124436&jump_to_nav=true) | The physical sexual characteristics of the neonate at birth. |
| Biospecimen Race | NCIT | [link](https://bioportal.bioontology.org/ontologies/NCIT/?p=classes&lang=en&conceptid=http%3A%2F%2Fncicb.nci.nih.gov%2Fxml%2Fowl%2FEVS%2FThesaurus.owl%23C92460&jump_to_nav=true) | A social environmental measure of neighborhood race and/or ethnic residential segregation based on data from the U.S. Census Bureau. |
| Treatment Dosage Regimen | NCIT | [link](https://bioportal.bioontology.org/ontologies/NCIT/?p=classes&lang=en&conceptid=http%3A%2F%2Fncicb.nci.nih.gov%2Fxml%2Fowl%2FEVS%2FThesaurus.owl%23C142516&jump_to_nav=true) | The specific way a therapeutic drug is to be taken, including formulation, route of administration, dose, dosing interval, and treatment duration. |

Abbreviations:

DEAM: The ontology of data analysis and management (1)

NCIT: National Cancer Institute Thesaurus (2)

ORNASEQ: Ontology of RNA Sequencing (3)

SO: Sequence Types and Features Ontology (4)

GFO-BIO: General Formal Ontology for Biology (5)

GEO: Gene Expression Omnibus

BCO: Biological Collections Ontology (6)

DOID: Disease Ontology (7)

OBI: Ontology for Biomedical Investigations (8)
