## Supplementary material for "RD-OMICS: An Integrative Multi-Omics Data Inventory in Rare Diseases": GEO_GBM_search.pdf

An official website of the United States government Here's how you know



**Entry type** clear
- ✓ **DataSets** (12)
- ✓ **Series** (814)
- Samples (0)
- Platforms (0)

**Organism**
Customize ...

**Study type**
Expression profiling by high throughput sequencing
Customize ...

**Author**
Customize ...

**Attribute name** clear
- ✓ **tissue** (826)
- strain (15)
- Customize ...

**Publication dates**
30 days
1 year
Custom range...

Clear all

Show additional filters

Summary ▾ 20 per page ▾ Sort by Default order ▾ Send to: ▾

### Search results

**Items: 1 to 20 of 826**

<< First < Prev Page 1 of 42 Next > Last >>

Filters activated: Series, DataSets, tissue. Clear all to show 34879 items.

1. Specific Intratumoral Microbiome Signatures in Human **Glioblastoma** and Meningioma: Evidence for a Gut–Brain Microbial Axis

   (Submitter supplied) Brain tumors (BTs), including **glioblastoma** (GBM) and meningioma (MGM), contribute significantly to the global cancer burden. The microbiome has been implicated in carcinogenesis, yet its role in BTs remains underexplored. We performed 16S rRNA gene sequencing of the gut microbiota (GM) and intratumoral microbiome (ItM) from fresh tissue samples of 9 patients with GBM and 18 with MGM. 12 age- and sex-matched healthy controls (HCs) were also enrolled. more...

   Organism: **Homo sapiens**
   Type: Other
   Platform: GPL15520 39 Samples
   Download data: CSV, FASTA
   Series Accession: GSE303995 ID: 200303995

2. Tumor-infiltrating lymphocytes-derived CD8+ clonotypes infiltrate the tumor tissue and mediate tumor regression in **glioblastoma** [WES]

   (Submitter supplied) IL-2/IL-15/IL-21-expanded TILs achieved complete tumor regression in a hypermutated **glioblastoma** patient, overcoming the immunosuppressive microenvironment. This case highlights a novel approach for treating gliomas using tailored adoptive cell therapy strategies.

   Organism: **Homo sapiens**
   Type: Other
   Platform: GPL24676 3 Samples
   Download data: TSV
   Series Accession: GSE285283 ID: 200285283
   PubMed Full text in PMC

**Filters:** Manage Filters

**Top Organisms** [Tree]
- Homo sapiens *(826)*
- Mus musculus *(53)*
- synthetic construct *(9)*
- Rattus norvegicus *(4)*
- Human betaherpesvirus 5 *(3)*
- More...

#### Find related data
Database: Select
Find items

#### Search details
```
("glioblastoma"[MeSH Terms] OR glioblastoma[All Fields]) AND "Homo sapiens"[porgn] AND (("gse"[Filter] OR "gds"[Filter]) AND "attribute name tissue"[Filter])
```
Search
See more...

#### Important Links
- GEO Home
- GEO Documentation
- About GEO DataSets

An official website of the United States government Here's how you know



Entry type
DataSets (0)
Series (17)
Samples (0)
Platforms (0)

Organism
Customize ...

Study type
Expression profiling by array
Methylation profiling by array
Customize ...

Author
Customize ...

Attribute name
tissue (8)
strain (1)
Customize ...

Publication dates
30 days
1 year
Custom range...

Clear all

Show additional filters

Summary 20 per page Sort by Default order Send to:

### Search results

**Items: 17**

The following term was not found in GEO DataSets: RNA-seq[Platform Technology Type].

See the search details.

1. Dual-targeting pharmacological UFMylation inhibition reprograms tumor and immune microenvironments to achieve long-term **glioblastoma** regression

(Submitter supplied) UFMylation, a recently identified ubiquitin-like modification, is essential for cellular stress homeostasis, particularly endoplasmic reticulum (ER) stress regulation. However, its biological and therapeutic exploration has been hindered by the absence of potent small-molecule inhibitors. Here, we report the first discovery of two compounds targeting the UFMylation E3 ligase complex core protein DDRGK1: Osimertinib, originally designed as an EGFR T790M selective inhibitor, acting through a previously unrecognized covalent mechanism, and CP-24, a novel non-covalent inhibitor. more...

Organism: **Homo sapiens**
Type: Expression profiling by high throughput sequencing
Platform: GPL34284 6 Samples
Download data: CSV
Series Accession: GSE280865 ID: 200280865

2. Cross-Species Transcriptomic Integration Reveals a Conserved, MIRO1-Mediated Macrophage-to-T Cell Signaling Axis Driving Immunosuppression in Glioma **[RNA-Seq]**

(Submitter supplied) We generated **bulk** RNA sequencing (**RNA-seq**) data from human glioma surgical resections treated ex vivo with a MIRO1-binding compound (MR3) or vehicle **control**. Fresh tumor specimens were processed and cultured under controlled conditions prior to RNA extraction and high-throughput transcriptomic profiling to characterize treatment-associated gene expression changes within the tumor microenvironment (TME). more...

Organism: **Homo sapiens**
Type: Expression profiling by high throughput sequencing
Platform: GPL34284 7 Samples
Download data: CSV, TABULAR
Series Accession: GSE322979 ID: 200322979
PubMed Full text in PMC Similar studies

**Filters:** Manage Filters

Top Organisms [Tree]
Homo sapiens *(17)*
Mus musculus *(1)*

#### Find related data

Database: Select

Find items

#### Search details

```
(((("glioblastoma"[MeSH Terms] OR
Glioblastoma[All Fields]) OR
("glioblastoma"[MeSH Terms] OR
Glioblastoma[All Fields])) AND
"Homo sapiens"[Organism]) AND
```

Search See more...

#### Important Links

GEO Home
GEO Documentation
About GEO DataSets
Construct a Query
Download Options

#### Recent activity

Turn Off Clear
